# Astrocytic activation of EMMPRIN contributes to their pathological phenotype in ALS

**DOI:** 10.1101/2025.02.23.639749

**Authors:** Gloria Nwamaka Edozie, Jasmine Bélanger, Shanna Pigeyre, Ariane Gosselin, Marion Boyer, Antoine G. Godin, Silvia Pozzi

## Abstract

Amyotrophic Lateral Sclerosis (ALS) is a fatal disease characterised by the degeneration of upper and lower motoneurons. Onset and progression of the disease are determined by both cell-autonomous neuronal dysfunctions and non-cell-autonomous factors, mainly due to activation of glial cells such as astrocytes and microglia.

The Extracellular Matrix Metalloproteinases INducer (EMMPRIN), a glycoprotein expressed by various cell types including neurons, is the major activator of matrix metalloproteinases (MMPs) synthesis and release. EMMPRIN activation can be induced by peptidyl-prolyl isomerase A (PPIA), a chaperone protein with cis/trans isomerase activity, that exhibits cytokine- and chemokine-like behaviour. Previous studies showed that PPIA is highly released in the cerebrospinal fluid (CSF) of ALS patients and animal models where, by activating EMMPRIN on motoneurons, induces neuronal death.

Here, we show that EMMPRIN is expressed also by astrocytes, suggesting this cell type as sensitive as motoneurons to PPIA-mediated EMMPRIN activation. We observed that that PPIA-mediated EMMPRIN activation prompt astrocytes toward a pro-inflammatory profile. Interestingly, we found that this pathogenic profile can be reverted by an anti-EMMPRIN antibody. Finally, we provide evidence that the activation of EMMPRIN is relevant for mutant SOD1 and TDP-43 conditions.

In conclusion, we demonstrate that EMMPRIN activation in ALS occurs also in astrocytes where it exacerbates their pathological phenotype possibly contributing to the progression of the disease. Furthermore, we suggest the potential use of an anti-EMMPRIN antibody to reduce astrocytic activation during the disease.

## Introduction

Amyotrophic Lateral Sclerosis (ALS) is a fatal neurodegenerative disease characterized by upper and lower motor neurons degeneration leading to muscle weakness, paralysis and eventually death (1). Ten percent of ALS cases are due to inherited mutations (familial ALS, fALS), while ninety percent are of unknown causes (sporadic ALS, sALS) (2). Different genes/proteins are implicated in ALS (3). Among them, SOD1 and TDP-43 are of particular interest, with most of the mutant SOD1 mouse models perfectly recapitulating the pathology observed in patients (4), and TDP-43 proteinopathy (mislocalization and aggregation of wild-type (WT) or mutant TDP-43) found in more than ninety-five percent of all cases of ALS (5).

ALS is a non-cell-autonomous disease. In the central nervous system, motoneurons are the exclusive site of first pathological events, contribute to glial cells activation as well as to disease onset and the early phases of the disease (6,7). Microglia activation initiate before motoneuronal loss and symptoms onset (8). While in the early phases of the disease microglia have a protective phenotype, they switch to a neurotoxic profile in the later phases (9), contributing to worsening disease progression and reducing survival (7). As per microglia, also astrocytes activate before onset of symptoms (10) and contribute to motoneuronal death by releasing toxic factors (11–13). Interestingly, in addition to being neurotoxic, astrocytes also contribute to microglial proliferation and to their neuroprotective-to-neurotoxic switch (14). Thus, factors that contribute to astrocytic activation, early in the disease, can impair their physiological communication with both motoneurons and microglia, and their study may uncover novel therapeutic targets.

The Extracellular Matrix Metalloprotease INducer (EMMPRIN), also known as Basigin or CD147, is a multi-functional glycoprotein expressed by different cellular types (15). As its name suggests, EMMPRIN is considered the main activator of the signal cascade leading to matrix metalloproteases (MMPs) synthesis and release (15). This protein is composed of two extracellular domains (ECI and ECII) at the N-terminal, and a transmembrane domain (TM), followed by a cytoplasmic tail (CT) at the C-terminal (16). It exists both as a membrane bound (MB) receptor and as a soluble molecule (16). Levels of EMMPRIN glycosylation are crucial for maturation, translocation to the membrane and further release as soluble EMMPRIN (15), as well as for the downstream pathways leading to MMPs production (15,17). Different studies have highlighted the role of EMMPRIN in pathological processes occurring in cancer, multiple sclerosis and spinal cord injury (15,18) but very less is known about EMMPRIN in ALS. Only one paper reported an increase of soluble EMMPRIN in ALS patients serum compared to healthy controls, and a correlation of EMMPRIN levels with disease severity (19).

EMMPRIN activation occurs via extracellular dimerization of the ECI (MB/MB or soluble/MB) (20–24) or by Peptidyl Prolyl isomerase A (PPIA)-mediated isomerisation of a peptidyl-prolyl residue in the ECII (25,26). PPIA is an intracellular chaperone (27) and a pro-inflammatory molecule that, by activating the NF-kB pathway (28–31), has cytokine and chemokine-like behaviours (32,33). Interestingly, elevated levels of extracellular PPIA have been observed in the cerebrospinal fluid (CSF) of sporadic ALS patients as well as in SOD1^G93A^ mice (34), a well-established mouse model of ALS (35). By binding to motoneuronal EMMPRIN, PPIA, secreted by astrocytes, induces their death *via* the stimulation of MMP-9 release (34), a metalloprotease linked to motoneuronal vulnerability in ALS (36–38), highlighting the detrimental role of EMMPRIN activation in motoneuronal degeneration.

Here, we further investigated the role of EMMPRIN activation in ALS and showed that this glycoprotein is necessary for PPIA-mediated NF-kB pathway. Moreover, we report that motoneurons are not the only cell type sensitive to PPIA. We found that astrocytes also express EMMPRIN and that in SOD1^G93A^ cells and mice the levels of this glycoprotein are increased. We evaluated the consequences of the astrocytic PPIA stimulation by investigating both the factors released and the relative activated pathways. We observed that PPIA-mediated EMMPRIN activation prompts astrocytes toward a pro-inflammatory profile, which can be reduced by a functional blocking antibody against EMMPRIN. Finally, we provide evidence that the activation of the pathway is relevant also in mutant TDP-43 conditions, extending our observations to non-SOD1-related ALS cases.

These findings underscore the crucial role of EMMPRIN in the activation of astrocytes during ALS, highlighting it as a promising therapeutic target across both SOD1- and TDP-43-related disease forms.

## Material and Methods

### Mice and Ethical Approval

All animal studies were approved by the Animal Care Ethics Committee of Université Laval under the protocol #2021-760 in accordance with the Guide to the Care and Use of Experimental Animals of the Canadian Council on Animal Care. Male transgenic SOD1^G93A^ mice were obtained from Jackson Lab (no. 004435) and maintained on a C57BL/6 background by breeding with wild type/non-transgenic (NTg) females (no. 000664). TDP-43^A315T^ mice (39–41) were provided by Dr. Jean-Pierre Julien. All mice were maintained and bred at animal facility of CERVO Brain Research Center under standard conditions: temperature 21 ± 1°C, relative humidity 55 ± 10%, 12h light schedule, food, and water *ad libitum*.

### Tissues collection

Tissue collection was performed following deep anesthesia of the mice with pentobarbital (12 mg/ml) at 0.01 ml/g. Cerebrospinal fluid (CSF) was collected from the cisterna magna using a capillary tube and centrifuged at 13 500 g for 5 min at 4°C. The supernatant was stored at −80°C until analysis. Following decapitation, the spinal cord was rapidly flushed out from the vertebral column, frozen on dry ice, and stored at –80°C until used for biochemical analyses. For immunofluorescence analysis, mice were perfused transcardially with cold 0.1M phosphate buffer (PB) followed by 4% paraformaldehyde (PFA) (Electron Microscopy Sciences, #19210) solution in PB 0.1M. Spinal cords were collected, post-fixed in 4% PFA for 24h at 4°C, then transferred in 30% sucrose in 0.1M PB.

### Primary astrocytes and treatment

Primary cultures of astrocytes were prepared from brain cortices of non-transgenic or SOD1^G93A^ mouse pups at 3-4 days old. Following removal of the meninges, the cerebral cortices were dissected and dissociated with 2 U/mL papain (Worthington) in Dulbecco’s Modified Eagle’s medium (DMEM) (Gibco, #12800-017) completed with cysteine and sodium bicarbonate at 37°C for 7 min. Tissues were dissociated with the addition of DNAse I (Sigma, #D5025) in the papain solution and incubated at 37°C for another 3 min. The papain was inactivated by subsequent washes of the cell suspension with DMEM supplemented with 10% fetal bovine serum (FBS) (Corning, #CA76322-116) and sodium carbonate. The clumps of cortices were further dissociated into single cells by trituration with needles (21G), then strained through a 70 μm cell strainer and centrifuged at 2 800 rpm for 5 min. Cells were resuspended and maintained in complete DMEM with 10% FBS and 1% penicillin/streptomycin (Gibco) and plated onto T-75 flasks at around 5×10^6^ cells. Cells were maintained at 37°C in a humidified 5% CO_2_ chamber until confluence, then shaken overnight at 200 rpm to remove microglia. Astrocytes were allowed to proliferate in culture for 14 days with the culture medium changed every 3-4 days. Astrocytes purity was confirmed by seeding cells on 10ng/mL poly-D-Lysine-coated coverslips for immunofluorescence analysis. Once confluence reached (day 15, DIV15), medium was changed and astrocytes were treated with 0.5nM recombinant mouse PPIA (Novus Biologicals, #NBP2-51988), 0.5 nM rat anti-EMMPRIN antibody (Invitrogen, #16-1471-82) or 0.5nM rat igG2A K isotype control (Invitrogen, #14-4321-82) in the culture media, or untreated. Cell lysates and conditioned medium were collected after 24 hours of treatment for biochemical analyses.

### Generation of the stable cell lines Hek-p65-luc, transfection and treatments

Human embryonic kidney cells (Hek-293) were cultured in DMEM supplemented with 10% FBS and 1% penicillin-streptomycin and were maintained at 37°C and 5% CO_2_. The NF-kB/luciferase reporter plasmid pGL4.32[*luc2P*/NF-κB-RE/Hygro] (Promega, #E859A) was stably transfected into Hek-293 cells using Jet Prime reagent (Polyplus), followed by selection and maintenance with hygromycin at 0,4 µg/mL. After reaching about 70% confluency, Hek-p65-luc cells were transfected with 0.5 µg of pcDNA3.1 (mock plasmid), human SOD1^WT^, SOD1^G93A^, TDP-43^WT^ or TDP-43^mNLS^ plasmids using Jet Prime reagent (Polyplus).

To further understand the involvement of EMMPRIN in PPIA-mediated NF-kB activation, small interference RNA (siRNA) was transfected into the cells to downregulate EMMPRIN expression. A mixture of four siRNA were used as a single reagent, providing advantages in both potency and specificity for EMMPRIN downregulation, using L-010737-00-0005-ON-TARGET plus SMARTpool for Human BSG, ((GGUCAGAGCUACAUUGA, GAAGUCGUCAGAACACAUC, GUACAAGAUCACUGACUCU, GGACAAGGCCCUCAUGAAC) (GE Healthcare DharmaconTM, Inc. Lafayette, CO, USA), or Non-Targeting-control-siRNA (UAGCGACUAAACACAUCAA, UAAGGCUAUGAAGAGAUAC, AUGAACGUGAAUUGCUCAA) (D-001810-10-05ON-TARGET plus Control Pool, non-targeting pool) (DharmaconTM) as control. 100 nmol/L of siRNA were transfected using JetPrime. Reduction of EMMPRIN was confirmed after 72h by western blot. Cells were then stimulated with 0.5nM recombinant mouse PPIA (Novus Biologicals) for 24h or co-treated with PPIA 0.5nM and anti-EMMPRIN antibody 0.5nM (Invitrogen,) or control antibody IgG2A isotype control 0.5nM (Invitrogen) for 24h. All treatments were performed in serum free media.

### Immunohistochemistry and immunocytochemistry

Primary cortical astrocytes (DIV15) were fixed with 4% PFA solution for 15 min at RT. Fixed cells were washed three times with phosphate buffered saline (PBS) 1X for 5 min each and then permeabilized in PBS containing 0.25% (v/v) Triton X-100 (PBS-T) for 15 min at RT. Cells were incubated in blocking solution containing 10 % normal goat serum (NGS) (Gibco, #16210072) in PBS-T for 1h, then incubated overnight at 4°C with anti-GFAP (astrocytes) and Iba-1 (microglia) primary antibodies diluted in blocking solution. Following washes, cells were incubated with appropriate Alexa Fluor-conjugated secondary antibodies (Life Technologies) diluted in blocking solution for 1h at RT. Nuclei were stained using Hoechst 33342 (Life Technologies, #H3570) at 1:1000 in PBS 1X for 1 min at RT. Images were captured with a Nikon A1R HD LSM Confocal microscope and NIS Element AR Software Vs 5.2.1 with a 25× objective at scale bar of 100 μm.

Coronal lumbar sections of 25 μm thickness were cut from the spinal cord of PFA-perfused mice using a microtome (LEICA, SM 2000R, Leica Biosystems, Germany). Antigen retrieval was performed by incubating sections in boiled 0.1M sodium citrate buffer pH-6.0 for 20 min. Sections were permeabilized in PBS-T for 15 min at RT, followed by a 1h incubation in blocking solution containing either 10% NGS or 5% (w/v) BSA (Bio-Basic) in PBS-T. Sections were incubated overnight at 4°C with the respective primary antibodies diluted in blocking solution. Following washes with PBS-T 1X, sections were incubated for 1h at RT with the respective Alexa Fluor-conjugated secondary antibodies diluted in blocking solution. Nuclei were stained using Hoechst 33342 (Life Technologies) at 1:1000 in PBS 1X for 1 min at RT. Images of the ventral horn gray matter were acquired using a Nikon A1R HD LSM confocal microscope and NIS Element AR Software Vs 5.2.1 with a 25× objective at scale bar of 100 μm. Image analysis was done using FIJI ImageJ (NIH).

To evaluate the percentage of cells (astrocytes or microglia) expressing EMMPRIN a colocalization analysis was performed on relative images. A macro was developed in ImageJ. Briefly, hyperstack images were loaded, Z-projection (Maximum intensity) obtained, and channels split. The ventral horn of the lumbar spinal cord was selected manually on the GFAP channel. All channels were thresholded, and the ventral horn selection was applied on each channel. Colocalization images between EMMPRIN and GFAP or EMMPRIN and Iba1 signals were generated using “logical AND operations” function in ImageJ. On GFAP or Iba1 channels a mask was created to obtain ROIs corresponding to individual cells. These ROIs were applied on the relative colocalization images and the percentage of cells expressing EMMPRIN was calculated and reported as fraction area. Threshold was determined for each channel using Auto Threshold and selecting the best fit (Default for EMMPRIN and Yen for Iba1 and GFAP).

### Protein extraction for tissues, cells and cellular media

Ventral horns of the lumbar spinal cord were obtained by longitudinal cryostat (CryoStar NX50, Thermo Fisher) separation from the dorsal horns. Total proteins from spinal cord or cells were lysed in 1% boiling sodium dodecyl sulfate (SDS). Briefly, samples were homogenized, sonicated, boiled 10 min at 99°C, and cleared by centrifugation at 12 000 rpm for 10 min. The supernatant fraction was collected and resuspended in loading buffer supplemented with dithiothreitol (DTT) (Fisher Scientific, # BP172-5). Proteins were quantified by DC protein assay (Bio-Rad, #50000112).

Cells media were collected and cleared from cell debris by centrifugation at 2 000 rpm for 5 min at 4°C. Supernatants were stored at –80°C for further analyses. For western blot, the supernatant fraction was precipitated with 4 volumes of cold acetone at –80°C overnight and pelleted by centrifugation at 15 000 *g* for 20 minutes. Proteins were resuspended in loading buffer supplemented with DTT.

### Dot blot and western blots

Dot Blot was performed by loading 2 μL of CSF on 0,45 µM PVDF membranes (Bio-Rad) by vacuum deposition on the Bio-Dot apparatus (Bio-Rad, # 1706545). Equal amount (30 μg) of proteins from cells or tissues were run on an 12% stain free acrylamide gel (Bio-Rad, # 1610185). After activation by UV light, gels were transferred onto 0,45 µM PVDF membranes (Bio-Rad). Ponceau S staining (0.1 % of dye in 5% of acetic acid) (Fisher Bioreagents) was performed to reveal total proteins loaded on the membrane after dot blots while Stain-Free UV-acquisition (41,42) of the membrane was performed for western blots. Membranes were blocked with 3% BSA (Bio-Basic) prepared in 0.1% Tween 20 (Fisher Scientific, # BP337-500) in Tris-buffered saline pH 7.5 (TBS-T) for 1h at RT, incubated with primary antibody overnight at 4°C, then incubated with appropriate peroxidase-conjugated secondary antibody for 1h at RT. Chemiluminescence blots were developed using Clarity™ Western ECL Substrate (Bio-Rad) on the ChemiDoc XRS imaging system (Bio-Rad). Densitometry was done with Image Lab software v6.1 (Bio-Rad). Immunoreactivity was quantified by Image Lab software v6.1 (Biorad) and normalized on total transferred proteins (TTP) obtained after Ponceau or stain free acquisition and quantification (Image Lab software).

### Dual Luciferase assay

Luciferase activity was measured using Bright-Glo Luciferase assay system (Promega, #E2620) according to the manufacturer’s protocol. Briefly, cells were lyses in 200 μL of 1X BrightGLO lysis buffer. Lysates were split in 25 µL triplicates in a 96-well plate, and 25 µL of luciferin, the luciferase substrate, was added to cell lysates. Relative luminescence units (RLU) were read using an EnSpire 2300 Multilabel reader (Perkin Elmer) at 490 nm. Total proteins were quantified from lysates using DC protein assay kit (Bio Rad). Relative luminescence units (RLU) were normalized on total protein for each sample.

### Cytokine array

Cytokine levels in astrocyte media were measured using a mouse cytokine array (C1000-Mouse antibody cytokine array, Ray Biotech, #AAM-CYT-1000-8) according to the manufacturer’s protocol. Medium (400 μL) from independent experiments were pooled and incubated with the antibody array membrane overnight at 4°C, after blocking the array with the proper blocking solution for 30 min. Membranes were incubated with biotinylated detection antibody overnight at 4°C before being incubated with streptavidin–conjugated secondary antibody at RT for 2 hr. Membrane was developed using ChemiDoc XRS imaging system by incubating equal mixture of array detection buffer 1 and 2. Experiments were run in technical duplicates. Densitometry was done using image Lab software v6.1 (Bio-Rad). Immunoreactivity values were normalized on the mean of the intensity of the positive control present on the different array membranes, as per manufacturer’s instructions.

### ClueGo analysis

Data from cytokine arrays were analyzed with ClueGo application (43,44) (version 2.5.9) using the Cytoscape environment(45) (version 3.9.1) as previously described (46). Differentially expressed proteins (with corresponding fold changes and p values) were used to generate biological networks using the following ontology sources: Gene Ontology (GO), Kyoto Encyclopedia of Genes and Genomes (KEGG), Reactome, and WikiPathways. The GO interval was between 3 (Min level) and 8 (Max level). The Kappa score was 0.5. For the enrichment of biological terms and groups, we used the two-sided (Enrichment/Depletion) tests based on the hyper-geometric distribution. We set the statistical significance to p<0.05, and we used the Bonferroni adjustment to correct the p value for the terms and the groups created by ClueGO. The leading group term is based on %genes/term vs. cluster.

### Statistical analyses

Means and standard deviation error (SEM) were used for all statistical analyses that were performed using GraphPad Prism 9 (GraphPad, San Diego, CA). The level of significance was specified as follows: * *p* < 0.05, ** *p* < 0.01, \*\*\**p* < 0.001 and **** *p* < 0.001. Unpaired T-test, One-way Anova or Two-way Anova followed by post hoc tests were performed depending on the experimental design as specified in figures’ legends.

### List of antibodies

Antibodies used for immunoblot, (western/dot blot) (IB), immunofluorescence (IF) are as follows: rat monoclonal anti-EMMPRIN antibody (1:1000 for IB; 1:500 for IF; Bio-Rad, #MCA2283), rabbit polyclonal anti-EMMPRIN antibody (1:1000 for IB; ProteinTech, #11989-1-AP), rabbit polyclonal anti-PPIA antibody (1:5000 for IB; ProteinTech, #10720-1-AP), rabbit polyclonal anti-NF-kB p65 subunit antibody (1:1000 for IB; Santa Cruz Biotechnology, #sc-8008), rabbit polyclonal anti-phospo-NF-kB p65 (Ser536) antibody (1:1000 for IB; Cell Signaling Technology, #3033), mouse monoclonal anti-phospo-NF-kB p65 (Ser536) antibody (1:1000 for IB; Cell Signaling Technology, #3036), rabbit polyclonal anti-Iba-1 antibody (1:500 for IF; Wako, #019-19741), mouse monoclonal anti-GFAP antibody (1:500 for IF; Cell Signaling Technology, #3670), mouse monoclonal anti-PPIA antibody (1:500 for IF; Invitrogen, #39-1100), rabbit monoclonal anti-NeuN antibody (1:500 for IF; Cell Signaling Technology, #12943), goat polyclonal anti-choline acetyltransferase antibody (1:500 for IF; Millipore, #AB114P), goat anti-mouse, anti-rabbit or anti-rat peroxidase-conjugated secondary antibodies (1:5000 for IB; Jackson Immunoresearch Lab), goat Alexa Fluor 647 or 555 or 488 anti-mouse or anti-rabbit or anti-rat or anti-goat fluorophore-conjugated secondary antibodies (1:500 for IF; Invitrogen).

## Results

### EMMPRIN is required for PPIA-induced NF-kB activation

Previous studies showed that the downstream effect of PPIA treatment is NF-kB activation (28–31). To confirm that EMMPRIN is necessary for PPIA-mediated NF-kB activation in ALS conditions we generated the Hek-p65-luc cell line. By having the luciferase gene cloned downstream of the p65/NF-kB responsive element, this cellular model allows fast and reliable measurements of NF-kB activation (40,47). Hek-p65-luc cells were transfected with plasmids expressing human SOD1 either in its wild-type form (SOD1^WT^) or carrying the familial ALS substitution Gly to Ala in position 93 (SOD1^G93A^). Cells were stimulated 24h with 0.5nM recombinant PPIA, a concentration able to induce SOD1^G93A^ motoneuronal death (34), and tested for NF-kB activation by luciferase assay. As shown in Fig. 1A, we confirmed that PPIA induces NF-kB activation specifically in SOD1^G93A^ cells.

**Figure 1.**
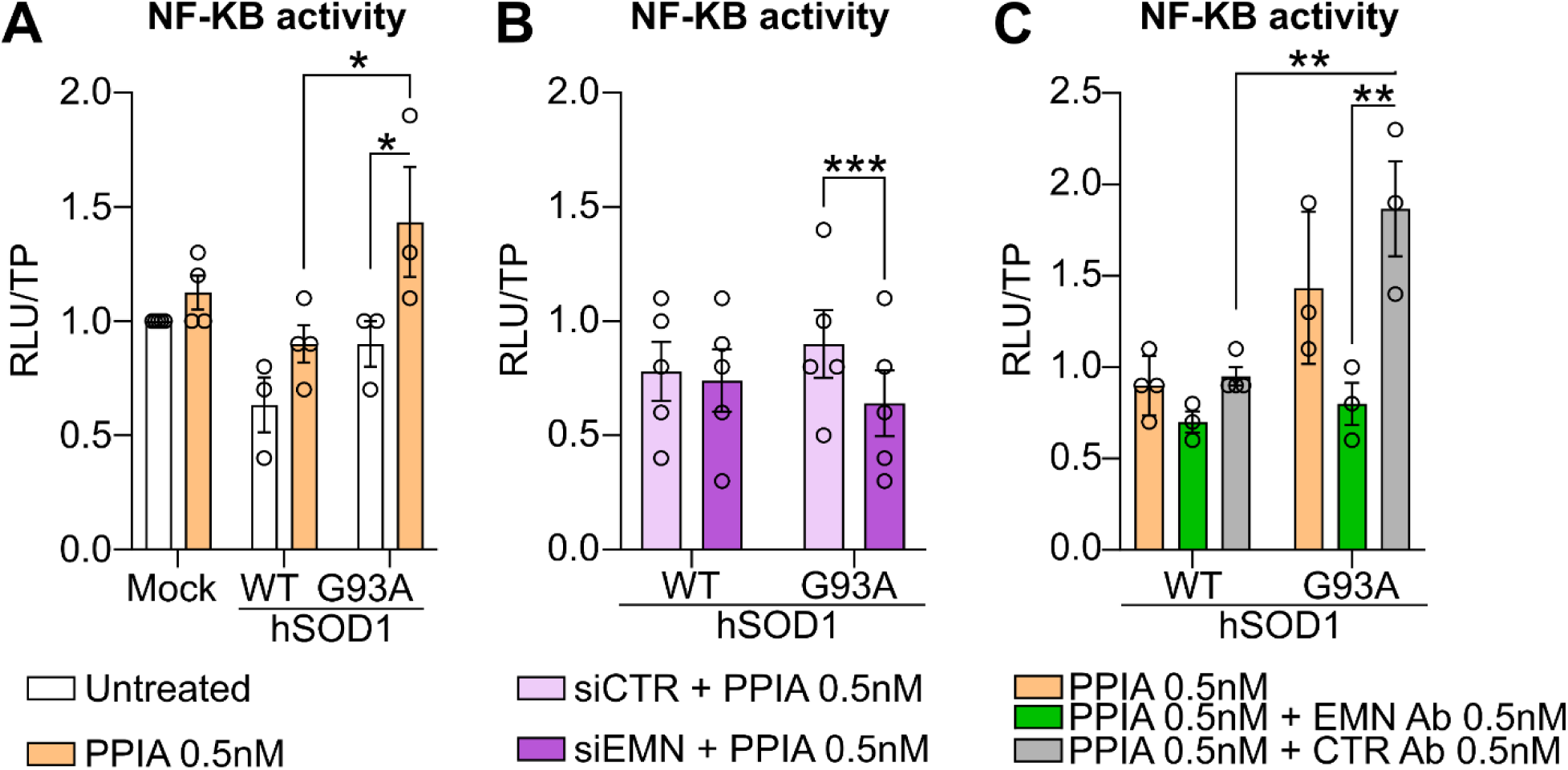
EMMPRIN is required for PPIA-mediated NF-kB activation in SOD1^G93A^ cells. **(A)** NF-kB activity in Hek-p65-luc cells transfected with human SOD1^WT^, SOD1^G93A^, or empty plasmid (mock) for 48h and treated with 0.5nM PPIA during the last 24h. Data are mean±SEM of n=3-4 independent experiments. Two-Way Anova followed by Tukey’s multiple comparisons test. **(B)** NF-kB activity in Hek-p65-luc cells transfected with siRNA control (siCTR) or against EMMPRIN (siEMN) for 72h, including 48h transfection with human SOD1^WT^ or SOD1^G93A^ and 24h treatment with 0.5nM PPIA. Data are mean±SEM of n=5 independent experiments. Two-Way Anova followed by Bonferroni’s multiple comparisons test. **(C)** NF-kB activity in Hek-p65-luc cells transfected with human SOD1^WT^ or SOD1^G93A^ plasmids for 48h and treated with a combination of 0.5nM PPIA and 0.5nM of control (CTR Ab) or anti-EMMPRIN (EMN Ab) antibody for the last 24h. Data are mean±SEM of n=3-4 independent experiments. Two-Way Anova followed by Tukey’s multiple comparisons test. For all experiments: All data were obtained by luciferase assay. Data are expressed as fold of Mock untreated cells. Relative luminescence units (RLU) were normalized on total proteins (TP, μg); *, p<0.05; **, p<0.01; ***, p<0.001.

We then investigated the role of EMMPRIN in PPIA-mediated NF-kB activation in SOD1^G93A^ cells by downregulating EMMPRIN using a small interference (siRNA) approach. Hek-p65-luc cells were first transfected with a control siRNA or a siRNA against EMMPRIN, then with plasmids expressing SOD1^WT^ or SOD1^G93A^ and lastly treated for 24h with 0.5nM recombinant PPIA. The luciferase assay revealed that the downregulation of EMMPRIN reduces PPIA-mediated NF-kB activation in SOD1^G93A^ cells (Fig. 1B).

Finally, to further confirm that NF-kB activation is induced by EMMPRIN activation in SOD1^G93A^ cells, we treated Hek-p65-luc cells with an anti-EMMPRIN functional blocking antibody, previously shown to block the PPIA-mediated EMMPRIN activation (48,49). Cells were transfected for 48h with plasmids expressing SOD1^WT^ or SOD1^G93A^ and treated with a combination, in the last 24h, of PPIA (0.5nM) and an anti-EMMPRIN antibody (0.5nM) or a control isotype antibody. As shown in Fig. 1C, the anti-EMMPRIN antibody treatment significantly reduces PPIA-mediated NF-kB activation in SOD1^G93A^ cells compared to the control antibody.

Overall, our results suggest that PPIA-induced NF-kB activation is EMMPRIN-dependent, and that, in presence of the familial ALS mutation SOD1^G93A^, cells are more sensitive to the PPIA-induced EMMPRIN stimulation.

### The PPIA/EMMPRIN pathway is more activated in SOD1^G93A^ cells

To further understand the sensitivity of SOD1^G93A^ cells to the PPIA-mediated EMMPRIN activation we analysed the protein levels of PPIA and EMMPRIN in Hek-p65-luc transfected cells by western blot (Fig. 2A,E). We observed that SOD1^G93A^ cells express more intracellular PPIA (Fig. 2B) as well as the low-glycosylated (Fig. 2C) and high-glycosylated (membrane bound) (Fig. 2D) forms of EMMPRIN. Furthermore, we observed that SOD1^G93A^ cells release more PPIA (Fig. 2F) as well as the soluble high-glycosylated form of EMMPRIN (Fig. 2G). These results suggest that in presence of SOD1^G93A^ mutation, cells are more sensitive to the PPIA-mediated activation of EMMPRIN due to an overexpression of these factors.

**Figure 2.**
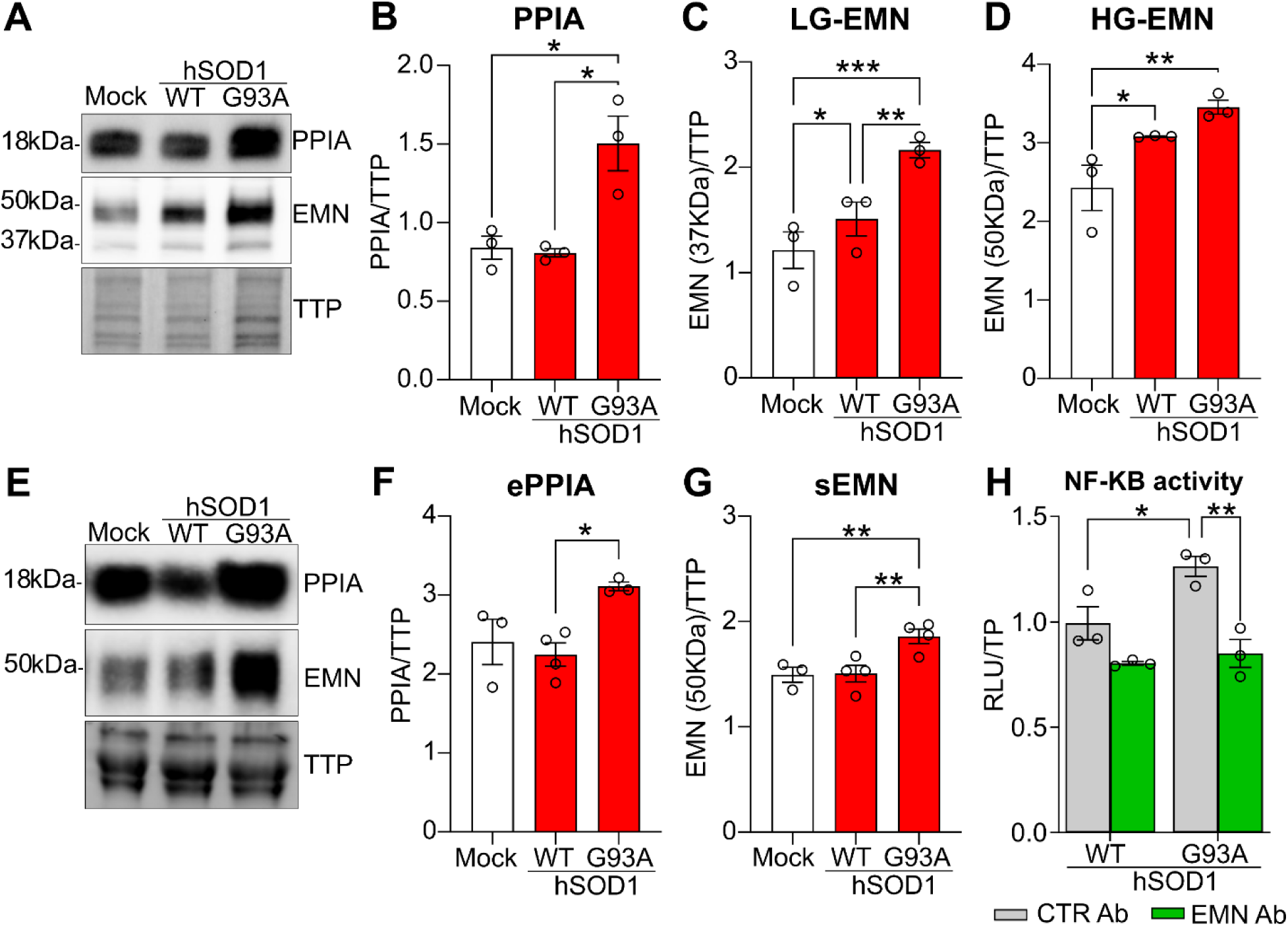
The PPIA/EMMPRIN pathway is more activated in SOD1^G93A^ cells. **(A-D)** Representative western blot of **(A)** lysates from 72h transfected Hek cells expressing human SOD1^WT^, SOD1^G93A^, or empty plasmid (mock) and relative quantification of PPIA **(B)**, low-glycosylated (37kDa) **(C)** and high-glycosylated (50kDa) **(D)** forms of EMMPRIN (EMN). Data are mean±SEM of n=3 independent experiments. One-Way Anova followed by uncorrected Fisher’s LSD test. **(E-G)** Representative western blot of **(E)** media from 72h transfected Hek cells expressing human SOD1^WT^, SOD1^G93A^, or empty plasmid (mock) and relative quantification of extracellular PPIA (ePPIA) **(F)** and soluble EMMPRIN (sEMN) **(G)**. Data are mean±SEM of n=3-4 independent experiments. One-Way Anova followed by uncorrected Fisher’s LSD test. **(H)** Luciferase assay for NF-kB activity in Hek-p65-luc cells transfected with human SOD1^WT^ or SOD1^G93A^ plasmids for 72h and treated with 0.5nM of control (CTR Ab) or anti-EMMPRIN (EMN Ab) antibody for the last 24h. Data are mean±SEM of n=3 independent experiments expressed as fold of Mock untreated cells. Two-Way Anova followed by Tukey’s multiple comparisons test. For all experiments: Target protein intensity was normalized on total transferred proteins (TTP). Relative luminescence units (RLU) were normalized on total proteins (TP, μg). *, p<0.05; **, p<0.01, ***, p<0.001.

Given the increased levels of EMMPRIN and extracellular PPIA in SOD1^G93A^ cells, we evaluated their NF-kB activation by luciferase assay. As shown in Fig. 1H SOD1^G93A^ cells have increased NF-kB activity compared to SOD1^WT^ transfected cells. This activation is interestingly reduced by a 24h treatment with the anti-EMMPRIN antibody suggesting that reducing EMMPRIN activation can modulate PPIA-mediated NF-kB activation in SOD1^G93A^ cells.

### EMMPRIN increases with disease progression in SOD1^G93A^ mice lumbar spinal cords, and it is expressed by astrocytes

To better understand the role of EMMPRIN in ALS pathology, we investigated its expression levels in SOD1^G93A^ mice with the disease progression. A western blot analysis was performed at different disease stages on the ventral horns of the lumbar spinal cord, where degenerating motoneurons and gliosis are present (Fig. 3A). Our results show a significant increase of the high-glycosylated form of EMMPRIN (membrane bound) in SOD1^G93A^ lumbar spinal cords already at the onset of symptoms, and a significant progressive increase with the disease progression (Fig. 3B). No changes in the expression of the low glycosylated form were detected in SOD1^G93A^ compared to NTg mice or as disease progresses (Fig. 3C) suggesting a constant recruitment of the newly produced EMMPRIN on the cellular membrane.

**Figure 3.**
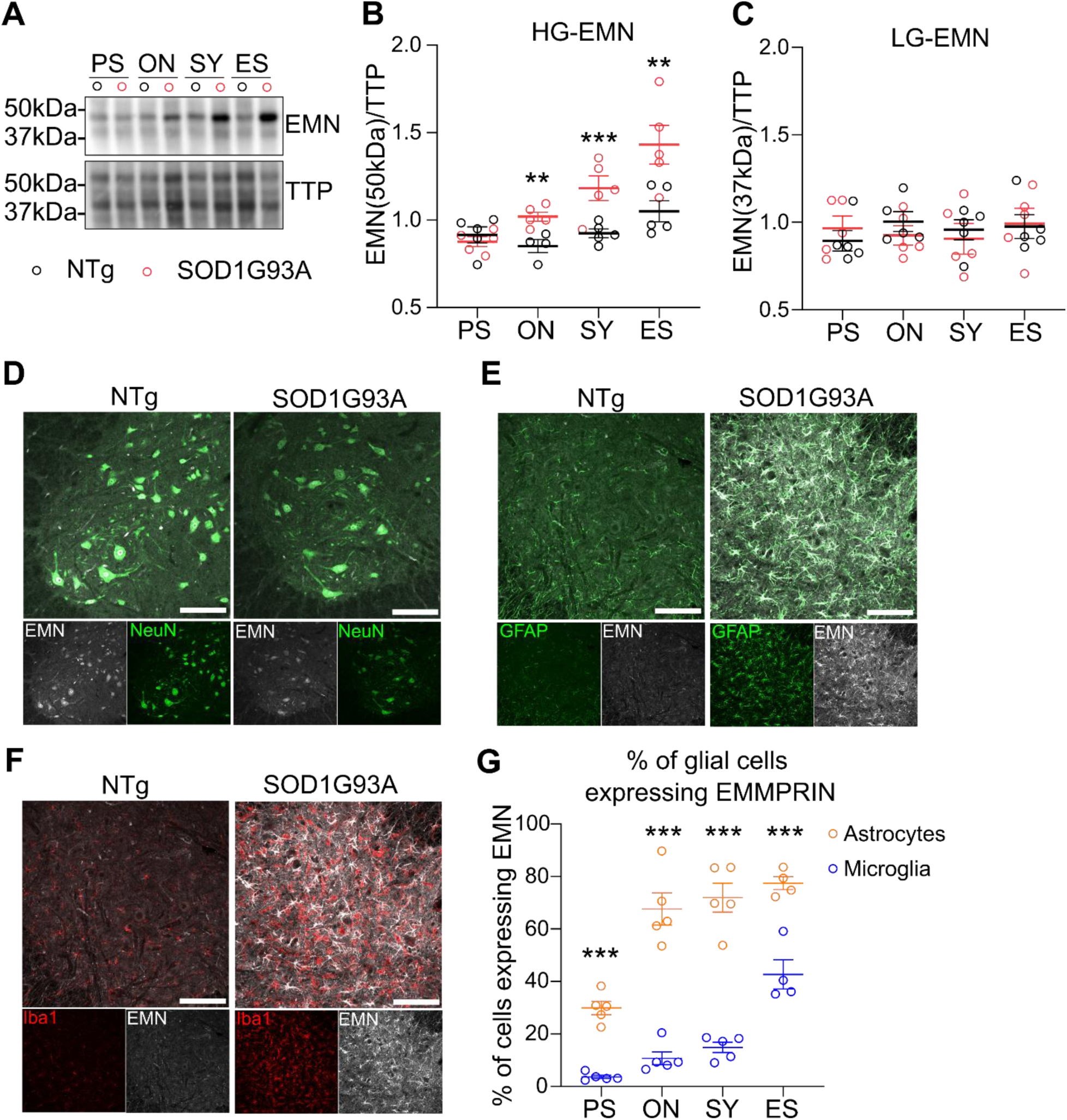
EMMPRIN increases with disease progression in SOD1^G93A^ mice, and it is expressed by astrocytes. **(A)** Representative western blot and relative quantification of the high-glycosylated (50kDa) **(B)** and the low-glycosylated (37kDa) **(C)** forms of EMMPRIN (EMN) in the ventral horns of the lumbar spinal cord of Ntg (black dots) and SOD1^G93A^ (red dots) mice. Two-Way Anova: HG-EMN (interaction, p=0.084; age, p<0.0001; genotype, p<0.0001) followed by T-test for genotype comparison: HG-EMN (PS, p=0.4986; ON, **, p=0.0064; SY, ***, p=0.0005; ES, **, p=0.0052); LG-EMN (interaction, p=0.6868; age, p=0.825; genotype, p=0.831). One-Way Anova for linear trend: HG-EMN, p<0.0001; LG-EMN, p=0.0636. Target protein intensity was normalized on total transferred proteins (TTP). Data are mean±SEM of n=5 mice/stage. **(D)** Representative image of EMMPRIN (EMN, gray) expression in neuronal cells (NeuN, green) in the ventral horn of the lumbar spinal cord of NTg and SOD1^G93A^ mice at the onset of the disease. Large neurons, i.e motoneurons, are labeled by anti-EMMPRIN antibody. Experiments have been performed in n=3 mice/group. **(E)** Representative image of EMMPRIN (EMN, gray) expression in astrocytes (GFAP, green) in the ventral horn of the lumbar spinal cord of NTg and SOD1^G93A^ mice at an advanced symptomatic stage. **(F)** Representative image of EMMPRIN (EMN, gray) expression in microglia (Iba1, red) in the ventral horn of the lumbar spinal cord of NTg and SOD1^G93A^ mice at an advanced symptomatic stage. Of note, E and F are the same image but with split channels and performed appropriate merge. **(G)** Relative quantification of the percentage of astrocytes (GFAP) or microglia (Iba1) expressing EMMPRIN. Two-Way Anova (interaction, p=0.0003; age, p<0.0001; cell type, p<0.0001) followed by Bonferroni multiple comparison test for cell type comparison (***PS, p=0.0001; ON, SY, ES ****, p<0.0001). One-Way Anova for linear trend: GFAP, p<0.0001; Iba1, p<0.0001. Data are mean±SEM of n=4-5 mice/stage. For all experiment: PS, presymptomatic; ON, onset; SY, symptomatic; ES, end-stage. For D-F scale bar = 100μm.

It has been shown that EMMPRIN is expressed by motoneurons in SOD1^G93A^ mice (34). Given the observed increase of EMMPRIN with the disease progression and the pathological degeneration and death of motoneurons in this animal model, we investigated the presence of EMMPRIN on other cell types of the lumbar spinal cord and focused on glial cells, known to be activated and to proliferative with the progression of the disease (50). First, we confirmed the neuronal expression of EMMPRIN in the lumbar spinal cord and its presence also on large neurons of the ventral horns, i.e. motoneurons, of both Ntg and SOD1^G93A^ mice (Fig. 3D). We then performed an immunofluorescence for glial cells, i.e. astrocytes (GFAP) and microglia (Iba1), in the lumbar spinal cord of advanced-stage SOD1^G93A^ and age-matched NTg mice (Fig. 3E,F) and analysed the percentage of glial cells expressing EMMPRIN in SOD1^G93A^ mice (Fig. 3G). In both genotypes we observed a predominant expression of EMMPRIN in astrocytes (Fig. 3E). In SOD1^G93A^ mice, a 30% of astrocytes express EMMPRIN already at the pre-symptomatic stage (Fig. 3G). This percentage increases with disease progression reaching a plateau around 70% already at the onset of the disease. In contrast, the presence of EMMPRIN in microglia was almost undetectable (Fig. 3F) and the percentage of microglia cells expressing EMMPRIN ranged below the 15% throughout the disease progression increasing to a 40% only at the end-stage (Fig. 3G). Of note, in SOD1^G93A^ mice a diffuse staining was observed also outside cells, suggesting the presence of soluble EMMPRIN trapped into the extracellular matrix.

Overall, we showed that in SOD1^G93A^ mice EMMPRIN is expressed early by astrocytes, making this cell type sensitive to the extracellular PPIA levels present in the CSF. Astrocytes, that become activated over time, might contribute to a general increase of EMMPRIN in the lumbar spinal cord with the progression of the disease.

### Activation of EMMPRIN induces a pro-inflammatory profile in astrocytes

To further evaluate the effect of PPIA-mediated EMMPRIN activation in astrocytes, we produced primary cortical astrocyte cultures from non-transgenic mice (Fig. 4A) and treated them for 24h with 0.5nM recombinant PPIA. By western blot we evaluated the levels of EMMPRIN (Fig. 4B, C), and surprisingly observed that PPIA stimulates its expression, possibly increasing their sensitivity to the PPIA/EMMPRIN pathway. Furthermore, to establish the effect of PPIA-mediated EMMPRIN activation in astrocytes we analysed the medium of 24h PPIA-stimulated cells by cytokine array (Fig. 4D and Supplementary Table1). Among the 95 factors tested, in the medium of PPIA treated cells we observed 43 factors upregulated (45.3%), 3 proteins downregulated (IL-4, Lymphotactin/XCL1 and SDF-1-alpha/CXCL12-alpha, 3.1%) and 49 unchanged proteins (51.6%) (Fig. 4E). To better characterize the molecular pattern of PPIA-activated astrocytes we run a pathway analysis of the 43 upregulated proteins and identified the leading terms by Cytoscape (45) and the ClueGO cluster analysis (43,44) (Fig. 4F and Supplementary Table 2). As expected, the most populated terms related to “cytokine-cytokine receptor binding”, “signal receptor binding” and “receptor ligand activity” supporting an enhancement of the astrocytic cellular communication. The most populated term highlighted by the analysis was “cytokine-cytokine receptor interaction” with 26 identified proteins. Among them we found pro-inflammatory factors such as IFN-gamma, IL-12, IL-15, IL-1 alpha and beta, IL-2, IL-6 and IL-7 as well as TNF-alpha and components of the tumor necrosis superfamilies of ligands (TNFSF) and receptors (TNFRSF) such as TNF receptor I and II, OPG (TNFRSF11B), GITR (TNFRSF18) and TROY (TNFRSF19). In addition, in this cluster, we identified chemokines and chemokines receptors such as MDC (CCL22), MIP-3-beta (CCL19), Eotaxin-2 (CCL24), MIP-1-alpha (CCL3), Rantes (CCL5), GM-CSF, KC (CXCL1), I-TAC (CXCL11), an MIP-2 (CXCL2). The proinflammatory profile of astrocytes treated with PPIA was also characterized by significant pathways related to “TNF signalling” and “response to TNF”, “extrinsic apoptotic pathways”, as well as “leukocytes migration” and “leukocytes differentiation”, “positive regulation of lymphocytes activation” and “lymphocytes differentiation”, “regulation of lymphocytes immunity” and “regulation of adaptive immune response”. It is worth mentioning that the pathway analysis identified also “matrix metalloproteases” and “NF-kB signalling pathway” as significant terms, confirming the downstream activation of NF-kB after the PPIA/EMMPRIN interaction and its contribution in astrocytic MMPs release. Finally, among the three downregulated proteins we found IL-4, a potent anti-inflammatory cytokine with demonstrated effects in maintaining microglia into an anti-inflammatory and protective state (9,51).

**Figure 4.**
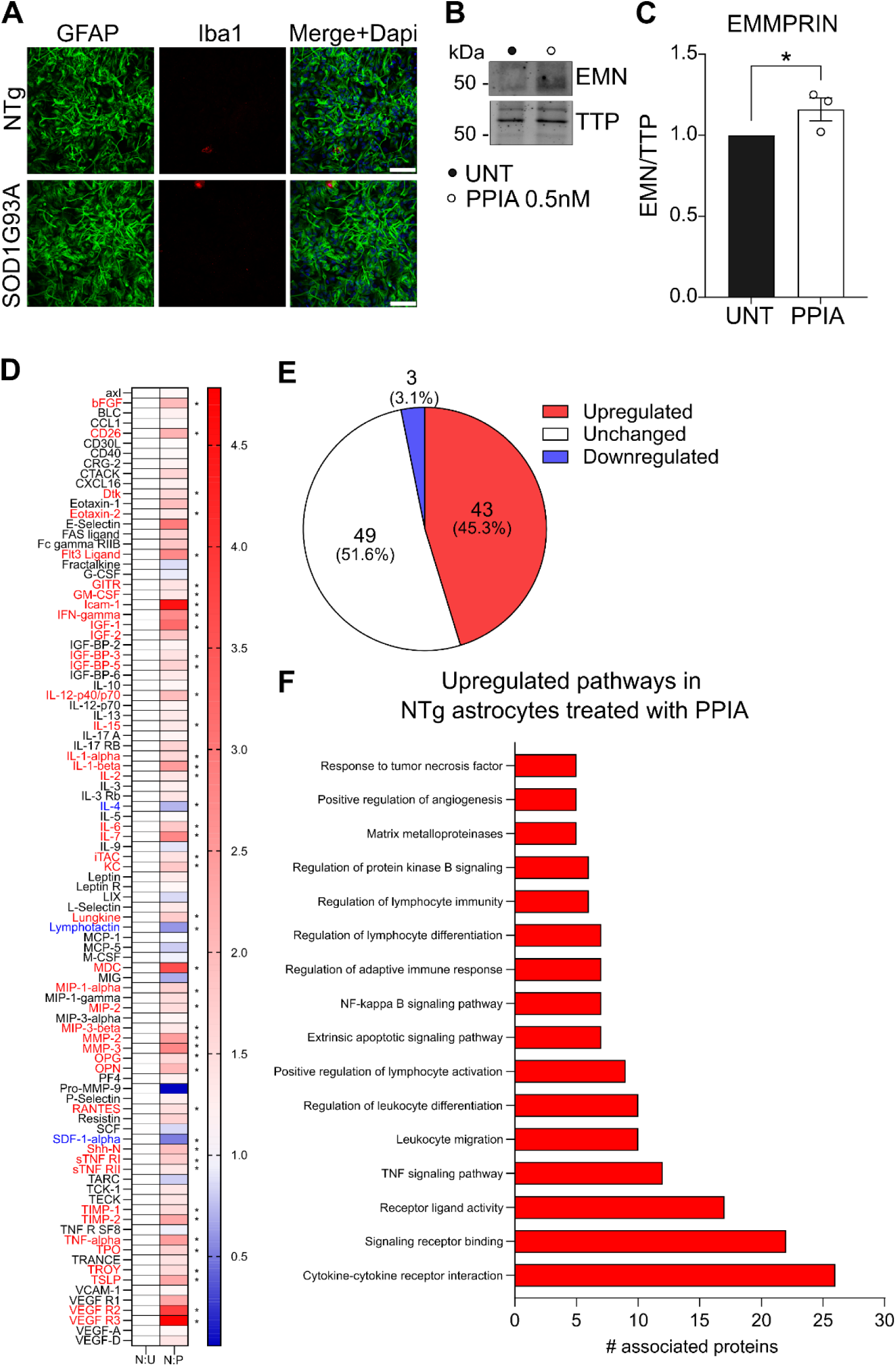
PPIA-mediated activation of EMMPRIN induces a pro-inflammatory phenotype in astrocytes. **(A)** Representative image of primary cultures of NTg and SOD1^G93A^ astrocytes. Most of the cells present in the preparation are astrocytes (GFAP), and only a very small percentage of microglia (Iba1) cells is detected. Scale bar = 100μm. **(B)** Representative western blot and **(C)** relative quantification of EMMPRIN (EMN) in NTg astrocytes treated 24h with 0.5nM recombinant PPIA. Data are mean±SEM of n=3 independent experiments (dots) expressed as fold of NTg cells. Target protein intensity was normalized on total transferred proteins (TTP). *, p<0.05 by unpaired T-Test. **(D)** Relative quantification of factors released by NTg astrocytes after 24h of treatment with 0.5nM recombinant PPIA (N:P) compared to untreated NTg astrocytes (N:U). Red (upregulated), black (unchanged), blue (downregulated). Pooled media of n=5 preparations in duplicate. Data are expressed as fold of NTg untreated cells (N:U). *, p<0.05 versus untreated NTg astrocytes by unpaired T-Test. Relative fold change, p-value, difference and q-value are listed in Supplementary Table 1. **(E)** Pie chart of upregulated, unchanged or downregulated proteins in NTg astrocytes treated with PPIA compared to untreated NTg astrocytes. **(F)** Significant leading pathways related to the 43 upregulated proteins found in NTg astrocytes treated with PPIA. Proteins associated with the pathways are listed in Supplementary Table 2.

Thus, PPIA-mediated EMMPRIN activation in astrocytes stimulates EMMPRIN expression, increasing their susceptibility to PPIA, and prompts them toward a pro-inflammatory profile.

### EMMPRIN, NF-kB and a pro-inflammatory profile are upregulated in SOD1^G93A^ astrocytes

To further understand the effect of EMMPRIN activation in mutant SOD1 conditions we produced primary cortical astrocyte cultures from SOD1^G93A^ mice (Fig. 4A) and investigated its expression and downstream activation by western blot (Fig. 5A-C). We confirmed previous observations showing that SOD1^G93A^ astrocytes release more PPIA than NTg cells (34) (Fig. 5A). Interestingly we found that SOD1^G93A^ astrocytes express more EMMPRIN compared to NTg astrocytes (Fig. 5B) which might be induced by the presence of PPIA in the medium. Because of PPIA-mediated EMMPRIN stimulation we also observed an increase of the phosphorylated form of p65 (Fig. 5C), the main subunit of NF-kB that becomes phosphorylated upon activation (40,52). These data suggest an autocrine activation of NF-kB mediated by the PPIA/EMMPRIN interaction in SOD1^G93A^ astrocytes. To better characterize the consequences of this activation in SOD1^G93A^ astrocytes, we ran a cytokine array on 24h conditioned medium of untreated NTg and SOD1^G93A^ astrocytes and observed an increase of different pro-inflammatory factors (Fig. 5D and Supplementary Table 3). Among the 95 factors analysed in cell media, we found 62 molecules (66%) upregulated, 31 proteins (33%) unchanged and only 1 protein downregulated (MCP-5, 1%) in SOD1^G93A^ astrocytes (Fig. 5E). Interestingly, when comparing the 43 upregulated factors after PPIA stimulation and the 62 factors upregulated in SOD1^G93A^ astrocytes, we found an overlap of 42 factors (Fig. 5F and Supplementary 4), most of them being pro-inflammatory factors and chemokines for immune cells recruitment. The pathway analysis on the 62 upregulated proteins found in the medium of SOD1^G93A^ astrocytes (Fig. 5G and Supplementary Table 5) revealed an intense astrocytic communication signalling characterized by a pro-inflammatory profile. Indeed, in the most populated term “cytokine-cytokine receptor interaction” we identified 39 proteins with pro-inflammatory and immune-modulating properties, such as IL-1 alpha and beta, IL-2, IL-6, IL-12 alpha and beta, IL-15, IL-17, IFN-gamma, TNF-alpha, components of the TNF superfamilies of ligands and receptors such as TNF RI and II, OPG (TNFRSF11B), GITR (TNFRSF18), TRACE (TNFSF11) and TROY (TNFRSF19), as well as chemokines and receptors such as MDC (CCL22), CCL1, MIP-3-beta (CCL19), Eotaxin-2 (CCL24), TECK (CCL25), CTACK (CCL27), MIP-1-alpha (CCL3), Rantes (CCL5), GM-CSF, IL-3-RB, KC (CXCL1), CRG-2 (CXCL10), I-TAC (CXCL11), CXCL16, MIP-2 (CXCL2), LIX (CXCL5) and MIG (CXCL9). The pro-inflammatory profile was supported also by the identification of other terms related to inflammation and inflammatory cells recruitment, such as “cytokine and inflammatory response”, “regulation of the inflammatory response”, “TNF-alpha signalling pathway” and “response to TNF”, “extrinsic apoptotic signalling pathways”, “IL-17 signalling pathway”, as well as pathways related to leukocytes and lymphocytes recruitment. Interestingly, 11 of the most significant pathways observed in SOD1^G93A^ astrocytes medium have a clear overlap with the analysis performed on the secretome of PPIA-treated NTg astrocytes (Fig. 5G, pathways with a star). Of note the 26 pro-inflammatory factors present in the “cytokine-cytokine receptor interaction” term of PPIA-stimulated NTg astrocytes were also found in the 39 factors that populated the same term in SOD1^G93A^ astrocytes. These observations highlighting the possible contribution of the PPIA-mediated EMMPRIN activation in the cellular response of ALS astrocytes. Moreover, among the common terms, we found also the “matrix metalloproteases” pathway, highlighting the presence of released MMPs, possibly after EMMPRIN activation, in SOD1^G93A^ astrocytic medium.

**Figure 5.**
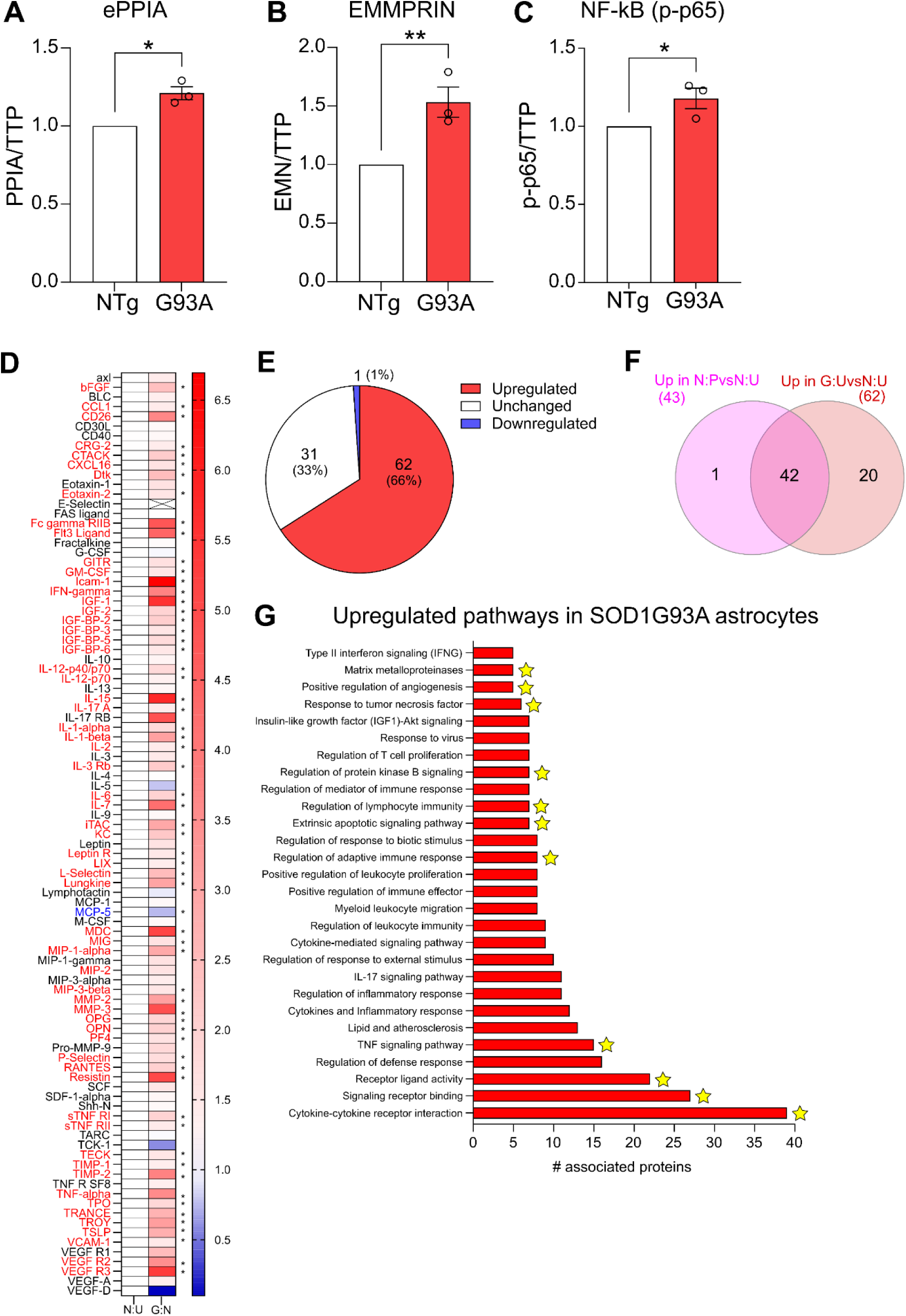
EMMPRIN, NF-kB and a pro-inflammatory profile are upregulated in SOD1^G93A^ astrocytes. Relative quantification of extracellular PPIA (ePPIA) **(A)**, EMMPRIN (EMN) **(B)** and the phosphorylated form pf p65/NF-kB (p-p65) **(C)** in NTg and SOD1^G93A^ astrocytes. Data are mean±SEM of n=3 independent experiments (dots) expressed as fold of NTg cells. Target protein intensity was normalized on total transferred proteins (TTP). *, p<0.05, **, p<0.01 by unpaired T-Test. **(D)** Relative quantification of factors released in 24h conditioned medium from SOD1^G93A^ astrocytes (G:U) compared to NTg astrocytes (N:U). Red (upregulated), black (unchanged), blue (downregulated). Pooled media of n=5 preparations in duplicate. Data are expressed as fold of Ntg untreated cells (N:U). *, p<0.05 versus untreated NTg astrocytes by unpaired T-Test. Relative fold change, p-value, difference and q-value are listed in Supplementary Table 3. **(E)** Pie chart of upregulated, unchanged or downregulated proteins in SOD1^G93A^ astrocytes compared to untreated NTg astrocytes. **(F)** Venn diagram showing 42 commonly secreted proteins between NTg astrocytes treated with PPIA (N:P) and SOD1^G93A^ astrocytes untreated (G:U). List of common proteins is present in Supplementary Table 4. **(G)** Significant leading pathways related to the 62 upregulated proteins found in SOD1^G93A^ astrocytes. Pathways found also in NTg astrocytes treated with PPIA are labelled with a star. Proteins associated with the pathways are listed in Supplementary Table 5.

Our results thus suggest that, in SOD1^G93A^ conditions, the activation of EMMPRIN, mediated by elevated levels of PPIA, may contribute to the pro-inflammatory profile of astrocytes that characterizes the disease (11,12,53).

### Blocking EMMPRIN activation reduces NF-kB activation and the pro-inflammatory profile of SOD1^G93A^ astrocytes

So far, we showed that PPIA-mediated activation of EMMPRIN induces the NF-kB pathway in SOD1^G93A^ transfected cells and NTg astrocytes treated with PPIA (Fig. 1, 2 and 4F). We also showed and that PPIA induces EMMPRIN expression (Fig. 4B, C), and that SOD1^G93A^ astrocytes release more PPIA, express more EMMPRIN, and show an increased NF-kB activation (Fig. 5A-C), suggesting an autocrine activation of the pathway in these cells. Finally, we observed an overlap between the secretome of SOD1^G93A^ astrocytes and NTg astrocytes treated with PPIA (Fig. 5F,G), suggesting the contribution of the PPIA/EMMPRIN pathway in the activation state of astrocytes in pathological conditions. To confirm this hypothesis, we treated SOD1^G93A^ primary astrocytes with the functional blocking antibody previously shown to reduce PPIA-mediated NF-kB activation in Hek cells (Fig. 1,2). First, we evaluated its effect on PPIA secretion, EMMPRIN expression and NF-kB activation by wester blot (Fig. 6A-C). We found that cells treated with anti-EMMPRIN antibody show a reduced secretion of extracellular PPIA (Fig. 6A) and expression of EMMPRIN (Fig. 6B) compared to control antibody treated cells. As a consequence of reducing the PPIA-mediated EMMPRIN activation we also observed a decrease of phosphorylated p65/NF-kB levels (Fig. 6C). We then investigated the efficacy of the anti-EMMPRIN antibody to reduce the pro-inflammatory phenotype previously observed in SOD1^G93A^ cells (Fig. 6D). The conditioned media of SOD1^G93A^ astrocytes treated for 24h with either a control isotype antibody or an anti-EMMPRIN antibody were assessed by cytokine array, with a particular focus on the 42 cytokines previously identified in common between SOD1^G93A^ astrocytes and NTg astrocytes treated with PPIA (Fig. 5F). As shown in Fig. 6D, E and Supplementary Table 6, the anti-EMMPRIN antibody significantly reduces the release of 30 factors (71.4%) previously found upregulated in non-transgenic cells treated with PPIA (Fig. 4D) and in SOD1^G93A^ astrocytes (Fig. 5D). 11 proteins (26.2%) were found unchanged in the secretome of SOD1^G93A^ treated with anti-EMMPRIN, whereas only 1 factor (IL-7, 2.4%) was upregulated after anti-EMMPRIN treatment. Interestingly, the pathway analysis run on the 30 downregulated protein after anti-EMMPRIN treatment (Fig. 6F and Supplementary Table 7), revealed a complete overlap with pathways found upregulated in SOD1^G93A^ astrocytes or NTg astrocytes treated with PPIA. Of note, in the most populated term of downregulated proteins after anti-EMMPRIN treatment, the “cytokine-cytokine receptor interaction”, we found the same pro-inflammatory cytokines or chemokines previously upregulated, such as IL-1-alpha and beta, IL-6, IL-12, TNF-alpha and components of the TNF family (OPG/TNFRSF11B, GITR/TNFRSF18, TNF RI and II), as well as MDC (CCL22), Eotaxin-2 (CCL24), Rantes (CCL5), I-TAC (CXCL11), KC (CXCL1) and MIP-2 (CXCL2). Moreover, although not evidenced by our pathway analysis, we also found a significant reduction of the levels of MMP-2 and MMP-3 released by SOD1^G93A^ astrocytes after anti-EMMPRIN treatment (Fig. 6D), which highlights a downregulation of matrix metalloproteases release after PPIA/EMMRPIN inactivation.

**Figure 6.**
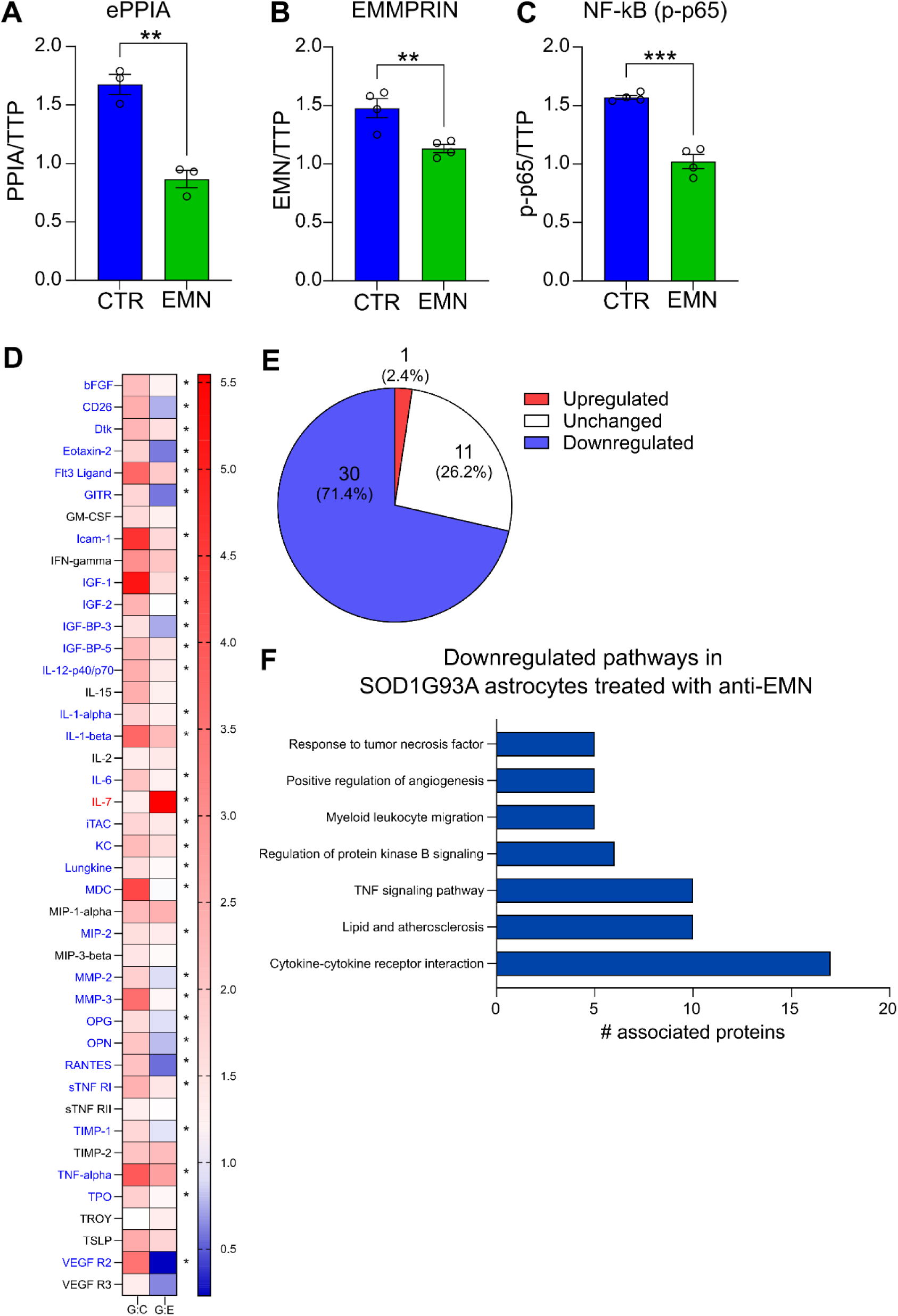
Blocking EMMPRIN activation reduces NF-kB activation and the pro-inflammatory profile of SOD1^G93A^ astrocytes. Relative quantification of extracellular PPIA (ePPIA) **(A)**, EMMPRIN (EMN) **(B)** and the phosphorylated form pf p65/NF-kB (p-p65) **(C)** in SOD1^G93A^ astrocytes treated 24h with 0.5nM anti-EMMPRIN (EMN Ab) or isotype control (CTR Ab) antibodies. Data are mean±SEM of n=3-4 independent experiments (dots) expressed as fold of NTg cells. Target protein intensity was normalized on total transferred proteins (TTP). *, p<0.05, **, p<0.01, ***, p<0.001 by unpaired T-Test. **(D)** Relative quantification of the 42 commonly secreted factors between untreated SOD1^G93A^ and NTg astrocytes treated with PPIA (List in Supplementary Table 4) released from SOD1^G93A^ astrocytes after 24h treatment with 0.5nM anti-EMMPRIN (G:E) or isotype control (G:C) antibodies. Red (upregulated), black (unchanged), blue (downregulated). Pooled media of n=4 preparations in duplicate. Data are expressed as fold of Ntg untreated cells (N:U). *, p<0.05; ** versus SOD1^G93A^ astrocytes treated with control antibody by unpaired T-Test. Relative fold change, p-value, difference and q-value are listed in Supplementary Table 6. **(E)** Pie chart of upregulated, unchanged or downregulated proteins in SOD1^G93A^ astrocytes treated with anti-EMMPRIN compared to control antibody treated SOD1^G93A^ astrocytes. **(F)** Significant leading pathways related to the 30 downregulated proteins found in SOD1^G93A^ astrocytes after anti-EMMPRIN treatment. Proteins associated with the pathways are listed in Supplementary Table 7.

Overall, our observations suggest that modulating EMMPRIN activation in SOD1^G93A^ astrocytes, with an anti-EMMPRIN functional blocking antibody, reduces PPIA release, EMMPRIN expression and NF-kB activation leading to a decrease of the proinflammatory phenotype of this cell population.

### The PPIA/EMMPRIN pathway is relevant also in mutant TDP-43 conditions

ALS is a disease with recognised 10% of cases due to familial inherited genetic alterations, and mutations in the human *SOD1* gene account only for 20% of familial cases (2,54). Thus, our results in SOD1^G93A^ conditions cover only a small proportion of all ALS forms. A hallmark of more than 95% of ALS cases (both sporadic and familial) is TDP-43 proteinopathy, described as the cytoplasmic mislocalization and aggregation of the protein TDP-43 (5). We therefore assessed the relevance of the PPIA/EMMPRIN interaction in mutant TDP-43 conditions.

To confirm that PPIA-mediates NF-kB activation also in TDP-43 conditions we transfected Hek-p65-luc cells with plasmids expressing human TDP-43^WT^ or TDP-43^mNLS^, a mutation that, by disrupting the nuclear localisation signal in TDP-43, mimics the cytoplasmic mislocalization and accumulation of TDP-43 observed during TDP-43 proteinopathy (41,55). As shown in Fig. 7A we observed a modest increase of NF-kB activity when cells express TDP-43^WT^, but a significant activation of NF-kB in cells expressing the mislocalized form of the protein. We then investigated the role of EMMPRIN in PPIA-mediated NF-kB activation by specifically downregulating EMMPRIN using a siRNA approach. Hek-p65-luc cells were first transfected with a control siRNA or a siRNA against EMMPRIN, then with plasmids expressing TDP-43^WT^ or TDP-43^mNLS^ and lastly treated for 24h with 0.5nM recombinant PPIA. The luciferase assay revealed that the downregulation of EMMPRIN in TDP-43^mNLS^ cells reduces PPIA-mediated NF-kB activation (Fig. 7B). Finally, we treated transfected Hek-p65-luc cells with the anti-EMMPRIN functional blocking antibody. As per the experiments with human SOD1, cells were transfected for 48h with plasmids expressing TDP-43^WT^ or TDP-43^mNLS^ and treated with a combination of recombinant PPIA (0.5nM) and an anti-EMMPRIN antibody (0.5nM) or control isotype antibody in the last 24h. As shown in Fig. 7C, the anti-EMMPRIN antibody treatment significantly reduces PPIA-mediated NF-kB activation both in TDP-43^WT^ and TDP-43^mNLS^ expressing cells.

**Figure 7.**
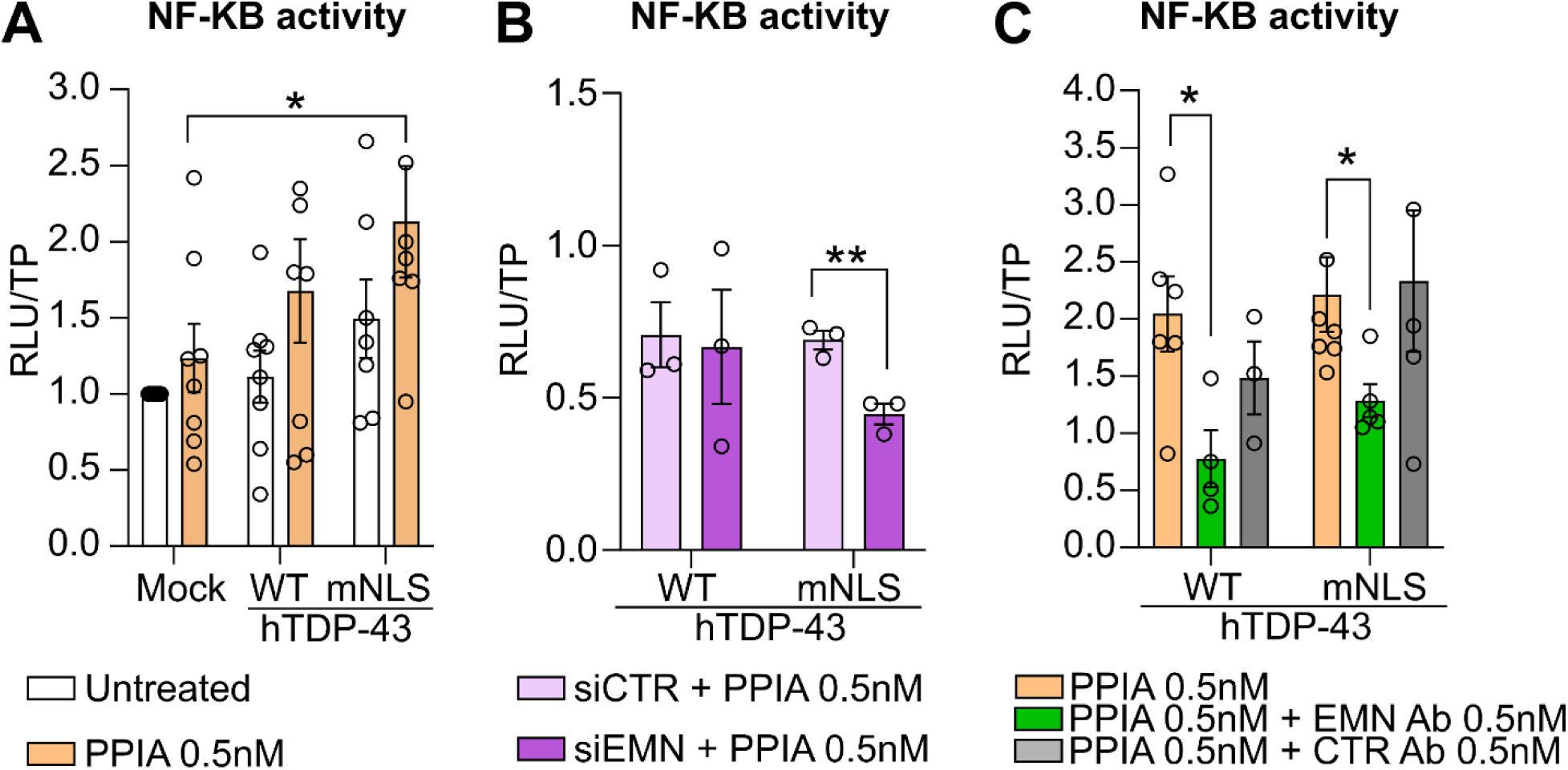
EMMPRIN is required for PPIA-mediated NF-kB activation in mutant TDP-43 cells. **(A)** NF-kB activity in Hek-p65-luc cells transfected with human TDP-43^WT^, TDP-43^A315T^, or empty plasmid (mock) for 48h and treated with 0.5nM PPIA during the last 24h. Data are mean±SEM of n=6-8 independent experiments. Two-Way Anova followed by Tukey’s multiple comparisons test. **(B)** NF-kB activity in Hek-p65-luc cells transfected with siRNA control (siCTR) or against EMMPRIN (siEMN) for 72h, including 48h transfection with human TDP-43^WT^ or TDP-43^A315T^ and 24h treatment with 0.5nM PPIA. Data are mean±SEM of n=3 independent experiments. Two-Way Anova followed by Bonferroni’s multiple comparisons test. **(C)** NF-kB activity in Hek-p65-luc cells transfected with human TDP-43^WT^ or TDP-43^A315T^ plasmids for 48h and treated with a combination of 0.5nM PPIA and 0.5nM of control (CTR Ab) or 0.5nM anti-EMMPRIN (EMN Ab) antibody for the last 24h. Data are mean±SEM of n=3-6 independent experiments. Two-Way Anova followed by Tukey’s multiple comparisons test. For all experiments: All data were obtained by luciferase assay. Data are expressed as fold of Mock untreated cells. Relative luminescence units (RLU) were normalized on total proteins (TP, μg); *, p<0.05; **, p<0.01; ***, p<0.001.

Overall, our results suggest that, in presence of TDP-43 mislocalization, cells are more sensitive to the PPIA-induced EMMPRIN stimulation.

To further investigate the pathway in TDP-43 conditions we analysed the levels of extracellular PPIA and EMMPRIN in a well-established mouse model of TDP-43 proteinopathy, the mouse expressing the familial mutation TDP-43^A315T^ (39–41). We first analysed the levels of extracellular PPIA in the CSF of these mice with the progression of the disease (Fig. 8A,B). As it occurs for SOD1^G93A^ mice, we observed an increase of extracellular PPIA at the onset of the disease. We then analysed the levels of EMMPRIN in the lumbar spinal cord of TDP-43^A315T^ mice by western blot (Fig. 8C) and observed a progressive increase of the high-glycosylated form of EMMPRIN with the progression of the disease (Fig. 8D) that correlated with a progressive decrease of the low-glycosylated form (Fig. 8E), suggesting an increase of the membrane-bound receptor with the disease. Finally, by immunofluorescence on lumbar spinal cord sections, we confirmed that EMMPRIN is expressed by neuronal cells and motoneurons (Fig. 8F), as well as by astrocytes (Fig. 8G) also in TDP-43^A315T^ mice.

**Figure 8.**
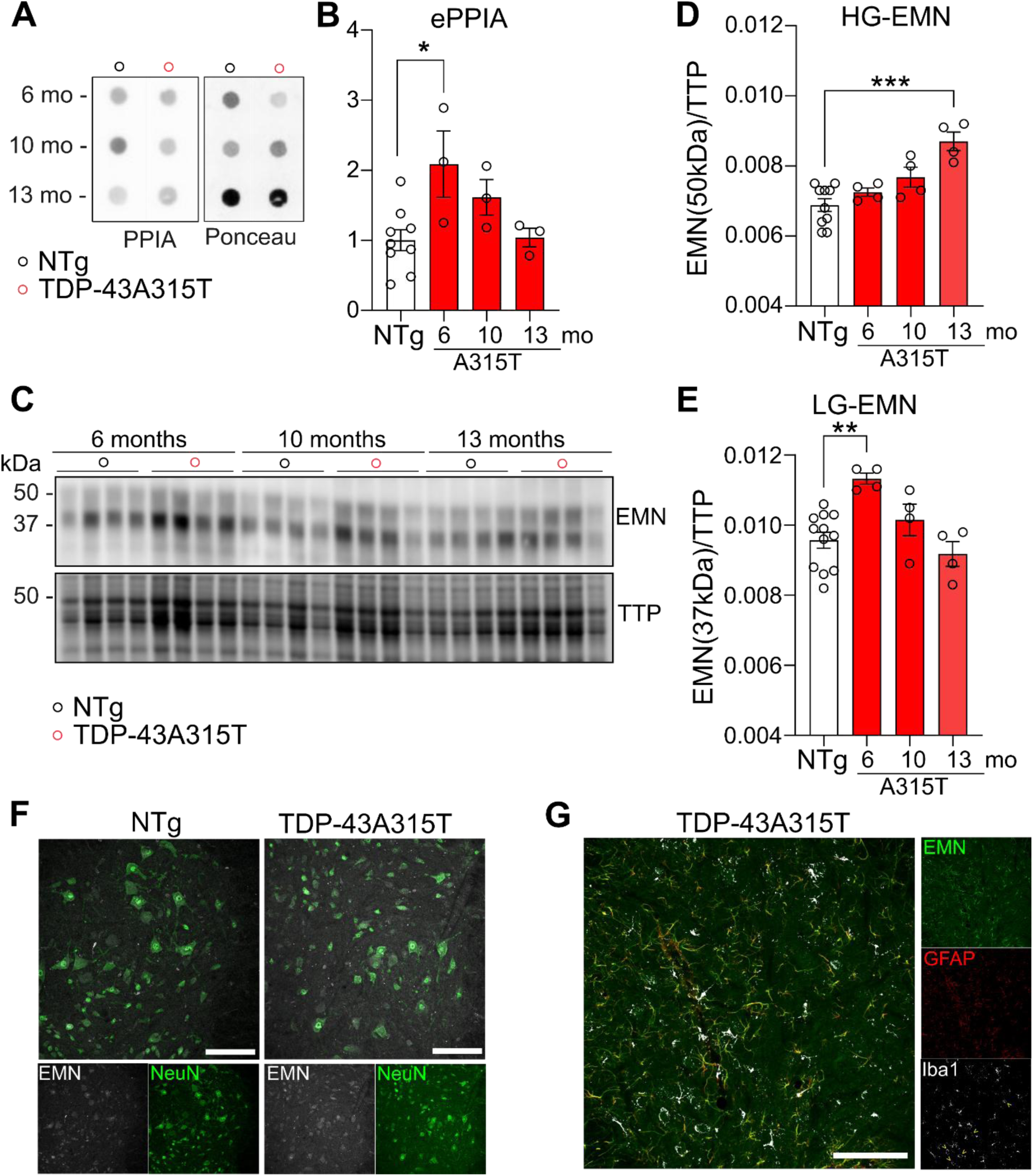
Extracellular PPIA and EMMPRIN are upregulated in TDP-43^A315T^ mice and EMMPRIN is expressed by astrocytes. Representative dot blot **(A)** and relative quantification (**B)** of extracellular PPIA in the CSF of Ntg and TDP-43^A315T^ mice. One-Way Anova followed by Tukey’s multiple comparison test. *, p=0.0248. One-Way Anova for linear trend in TDP-43^A315T^ mice: p=0.1466. Representative western blot (**C**) and relative quantification of the high-glycosylated (50kDa) **(D)** and the low-glycosylated (37kDa) **(E)** forms of EMMPRIN (EMN) in the lumbar spinal cord of Ntg and TDP-43^A315T^ mice. One-Way Anova followed by Tukey’s multiple comparison test. **, p=0.0020; ***, p=0.0001. One-Way Anova for linear trend in TDP-43^A315T^ mice: HG-EMN, p=0.0051; LG-EMN, p=0.0054. For A-E, target protein intensity was normalized on total transferred proteins (TTP). Data are mean±SEM of n=3-4 mice/stage (6 months, onset; 10 months, early symptomatic; 13 months, late-symptomatic). **(F)** Representative image of EMMPRIN (EMN, gray) expression in neuronal cells (NeuN, green) in the ventral horn of the lumbar spinal cord of NTg and TDP-43^A315T^ mice at the onset of the disease (6 months). Large neurons, i.e motoneurons, are labeled by anti-EMMPRIN antibody. **(G)** Representative image of EMMPRIN (EMN, green) expression in astrocytes (GFAP, red) or microglia (Iba1, gray) in the ventral horn of the lumbar spinal cord of TDP-43^A315T^ mice at the early symptomatic stage (10 months). Please note that due to antigen retrieval we observed some Iba1 leaking signal in neurons (yellow arrow heads). For F, G: experiments have been performed in n=3 mice/group. Scale bar = 100μm.

## Discussion

Extracellular PPIA is highly released in the CSF of SOD1^G93A^ mice and rats as well as in sporadic ALS patients (34). In this report, we demonstrate an increase of extracellular PPIA also in the CFS of TDP-43^A315T^ mice (Fig. 8) corroborating the evidence of an abundance of extracellular PPIA in ALS conditions. Which is the effect of extracellular PPIA in ALS? Previous studies have highlighted the toxic effect of the PPIA/EMMPRIN interaction on SOD1^G93A^ motoneurons (34). It has been demonstrated, indeed, that in primary motoneuronal cultures PPIA induces SOD1^G93A^ motoneuronal death by increasing MMP-9 expression, a matrix metalloprotease particularly toxic for ALS motoneurons (36–38). SOD1^G93A^ astrocytes release high levels of PPIA and a treatment with an extracellular inhibitor of PPIA reduces motoneuronal death in astrocytes-neurons co-cultures (34), highlighting the toxic role of a paracrine activation of motoneuronal EMMPRIN, mediated by the astrocytic release of PPIA. In this paper we show, for the first time, that in ALS, notably SOD1^G93A^ and TDP-43^A315T^ mice, EMMPRIN is expressed not only by motoneurons but also by astrocytes (Fig. 3, 8), suggesting a double effect of extracellular PPIA on both cell types during the disease.

PPIA is a chaperone protein (27) with extracellular cytokine and chemokine-like behaviour (32,33). By binding to EMMPRIN, PPIA induces its isomerization and the downstream upregulation of the EMMPRIN intracellular signal cascade (25,26,56). Here, by gene expression modulation and pharmacological intervention, we confirm that the downstream intracellular effector of this interaction is NF-kB and highlight the exclusive need of EMMPRIN for PPIA-mediated NF-kB activation (Fig. 1, 2, 4, 6, 7). Interestingly, we demonstrate that, in SOD1^G93A^ and mutant TDP-43 expressing cells, the NF-kB activation mediated by PPIA is higher than in control conditions, suggesting that ALS cells are more prone to respond to PPIA (Fig. 1, 7). This might be explained by the increase of EMMPRIN expression that we observed in SOD1^G93A^ cells (Fig. 2) and astrocytes (Fig. 5) as well as in SOD1^G93A^ and TDP-43^A315T^ tissues (Fig. 3, 8). Given the progressive degeneration of motoneurons, the specific localization of EMMPRIN in astrocytes over microglial cells, and the increased expression of EMMPRIN in SOD1^G93A^ astrocytes, we suggest that the general increase observed in mice tissues might be due to the astrocytic activation reported during the disease, highlighting astrocytes as sensitive cell type for EMMPRIN activation throughout the course of the disease.

Our observations in SOD1^G93A^ transfected cells, clearly suggested the need for PPIA/EMMPRIN interaction in NF-kB activation, indeed by reducing EMMPRIN levels with an siRNA approach, we lost the possibility for an interaction and observed reduced NF-kB activation (Fig. 1, 7). Of note, when treating SOD1^G93A^ astrocytes with an anti-EMMPRIN antibody we observed also reduced levels of EMMPRIN expression (Fig. 6B). It is known that antibodies can penetrate cells an reduce the intracellular levels of their target (41,57,58), therefore we could hypothesize the same effect of the anti-EMMPRIN antibody in our cellular model. Interestingly though, when reducing EMMPRIN levels with an siRNA approach we observed a 60% reduction in PPIA-mediated NF-kB activation (SOD1^G93A^ cells + PPIA compared to SOD1^G93A^ cells + PPIA + siEMN) whereas when treating with an anti-EMMPRIN antibody we observed a 47% reduction (SOD1^G93A^ cells + PPIA compared to SOD1^G93A^ cells + PPIA + anti-EMN) (Fig. 1), suggesting an exclusive extracellular effect of the antibody in our experimental conditions, and a possible regulation of EMMPRIN expression mediated by NF-kB activation. Of note, this is observable also in mutant TDP-43 conditions (Fig. 7) where the siRNA approach reduces by 75% PPIA-mediated NF-kB activation, whereas the anti-EMMPRIN antibody mediates a 35% reduction compared to mutant TDP-43 transfected cells treated with PPIA.

NF-kB is a transcription factor considered as a master regulator of inflammation (59,60). Different studies have reported the implications of NF-kB activation also in ALS (14,61,62) highlighting its specific and detrimental effects when activated in neurons (61), microglia (62) or astrocytes (14). In this paper, we report the activation of EMMPRIN as a new mechanism by which NF-kB is induced in ALS astrocytes. By treating healthy astrocytes with PPIA we highlight that PPIA-mediated activation of EMMPRIN prompt them toward an inflammatory phenotype characterized by the release of pro-inflammatory cytokines such IFN-gamma, IL-12, IL-15, IL-1 alpha and beta, IL-2, IL-6, and IL-7, as well as TNF-alpha and components of the tumor necrosis superfamilies, and chemokines such as MDC, MIP-3-beta, Eotaxin-2, MIP-1-alpha, Rantes, GM-CSF, KC, I-TAC, an MIP-2 (Fig. 4). Interestingly, we found an upregulation of the same factors in SOD1^G93A^ untreated astrocytes (Fig. 5), where we reported an increase of extracellular PPIA, as well as an increase of EMMPRIN expression and NF-kB activation. The inhibition of EMMPRIN activation, mediated by a functional blocking antibody against EMMPRIN, reduces NF-kB activation as well as the pro-inflammatory profile in SOD1^G93A^ astrocytes, suggesting the role of the PPIA/EMMPRIN interaction in the astrocytic activation during ALS, as well as an autocrine activation of the pathway in this cell type (Fig. 6). Furthermore, most of these factors are known to be upregulated in ALS, secreted by ALS activated astrocytes and active contributors of the pathology (53). Pro-inflammatory cytokines apart, our data also highlight an increased secretion of chemokines known to be involved in macrophages/microglia as well as lymphocytes recruitment in ALS (63). Finally, we also observed an increase of MMP-2 and MMP-3 in PPIA- stimulated and SOD1^G93A^ astrocytes media which is reverted by the anti-EMMPRIN antibody. Given the toxicity of MMP-9 for motoneuron, this observation is of particular interest, as it suggests that PPIA/EMMPRIN mediated interaction could contribute to the pathology *via* the release of different and cell-specific MMPs. Of note, highest levels of extracellular PPIA and EMMPRIN are observed at the onset of disease both in SOD1^G93A^ (34) and TDP-43^A315T^ mice (Fig. 3, 8), highlighting this stage of the disease as a peculiar window for the PPIA/EMMPRIN interaction. Interestingly, astrocytic activation of NF-kB at the onset of symptoms in SOD1^G93A^ mice appear to have a direct effect on microglia proliferation and pro-inflammatory/neurotoxic switch (14). We can therefore assume that, at this stage, a paracrine effect of astrocytes-released PPIA contributes to motoneuronal death (34), whereas an autocrine activation of the pathway in astrocytes prompt them toward a pro-inflammatory profile that might contribute to microglial proliferation and neurotoxic activation (14).

Our data show that SOD1^G93A^ transfected cells not only express high levels of EMMPRIN but also release its soluble form (Fig. 2), which may suggest an increased release of soluble EMMPRIN also by astrocytes. This might explain the presence of a diffuse staining observed in SOD1^G93A^ mice spinal cord at the advanced stages of the disease (Fig. 3). Of note, soluble EMMPRIN can activate its membrane bound form (20–24) suggesting a role for soluble EMMPRIN (under evaluations in the lab) in the advanced stages of the disease. This might suggest a constant activation of EMMPRIN during the disease progression with high levels of PPIA in the early phases that enhance, by isomerization, EMMPRIN activation, and a progressive increase of soluble EMMPRIN/membrane bound EMMPRIN activation throughout the disease progression, even with low levels of extracellular PPIA.

Finally, our observations suggest EMMPRIN as a possible target for therapeutic approaches in ALS and highlight the benefits of using an anti-EMMPRIN antibody to reduce its activation state and downstream consequences. Previous studies demonstrated that specific inhibition of the extracellular PPIA improves SOD1^G93A^ mice survival, preserves motoneurons from death and reduces astrogliosis (34). Although promising, this molecule is an analog of cyclosporine A (64,65) and might exhibit the same dose dependant toxicity of its precursor (66) in clinical applications, limiting the translational potential to patients. On the other side, there is nowadays an increasing interest in the use of monoclonal antibodies to treat neurodegenerative disorders (67,68), including ALS (69), and the antibody-based approach investigated in this study might be a novel therapeutic avenue (a preclinical trial is currently ongoing in our lab).

## Conclusions

Based on the obtained results, our study provides a novel understanding of EMMPRIN in ALS. Activation of EMMPRIN does not only occur in motoneurons but also in astrocytes during the disease and, in these cells, it contributes to their pathological profile. Furthermore, our observations prove that EMMPRIN activation is not restricted to SOD1 mutant forms, but it is relevant also in conditions with TDP-43 proteinopathy, observed in more than 95% of ALS patients. We thus conclude that in ALS, EMMPRIN activation has a dual role in the pathological processes, inducing both motoneurons degeneration and astrocytic activation, further contributing to the progression of the disease. The molecular aspects of these double effect though need to be still investigated. Finally, targeting EMMPRIN, with a specific anti-EMMPRIN antibody as suggested in this paper, may reveal as a promising therapeutic approach for ALS.

## Author contributions

S.P. conceived the study. G.E. performed most of the experiments and data analyses supported by J.B., Sh.P., A.G. and M.B. S.P. performed the pathways analyses. A.G.G. developed tools for image analysis. G.E. drafted the first version of the manuscript. S.P. revised and finalised the manuscript. All authors read and approved the manuscript.

## Acknowledgement and Funding

We thank Dr. Julien for providing the TDP-43^A315T^ mice and the plasmids expressing human SOD1^WT^, SOD1^G93A^, TDP-43^WT^ and TDP-43^mNLS^; Dr. Kriz for providing the SOD1^G93A^ mice for the first immunofluorescence analyses; Dr. Bonetto and Dr. Pasetto for first discussions.

This study was supported by the ALS Canada/Fondation Vincent Bourque Career Transition Award, the IBRO-USCRC Rising Star Award, and the Natural Science and Engineering Research Council of Canada (RGPIN-2021-03394) to S.P. who is a CIHR Tier 2 Canada Research Chair in Central and Peripheral Cellular Networks in ALS.

A.G.G. is a Junior 2 Scholar of the Fonds de recherche du Québec– Santé and was supported by a Sentinel North Partnership Research Chair and grant #06507 from the Natural Sciences and Engineering Research Council of Canada.

