## Supplementary material for "Astrocytic activation of EMMPRIN contributes to their pathological phenotype in ALS"

**Supplementary Table 1**

| NTg untreated (N:U) vs NTg + PPIA (N:P) | | | | | | | |
|---|---|---|---|---|---|---|---|
| | | Mean | | | SE of | | |
| **Factors** | **P value** | **N:U** | **N:P** | **Difference** | **difference** | **q value** | **in N:P** |
| axl | 0.6122 | 1.00 | 1.09 | -0.09 | 0.17 | 0.4975 | UNCHANGED |
| bFGF | 0.0000 | 1.00 | 2.00 | -1.00 | 0.06 | 0.0001 | UP |
| BLC | 0.1681 | 1.00 | 1.19 | -0.19 | 0.12 | 0.1748 | UNCHANGED |
| CCL1 | 0.5545 | 1.00 | 1.07 | -0.07 | 0.11 | 0.4580 | UNCHANGED |
| CD26 | 0.0148 | 1.00 | 2.12 | -1.12 | 0.33 | 0.0262 | UP |
| CD30/TNFRSF8 | 0.7103 | 1.00 | 0.95 | 0.05 | 0.13 | 0.5518 | UNCHANGED |
| CD30L | 0.6230 | 1.00 | 1.06 | -0.06 | 0.11 | 0.5005 | UNCHANGED |
| CD40 | 0.7593 | 1.00 | 1.08 | -0.08 | 0.23 | 0.5664 | UNCHANGED |
| CRG-2 | 0.2790 | 1.00 | 1.18 | -0.18 | 0.15 | 0.2692 | UNCHANGED |
| CTACK | 0.1517 | 1.00 | 1.61 | -0.61 | 0.37 | 0.1650 | UNCHANGED |
| CXCL16 | 0.2592 | 1.00 | 1.20 | -0.20 | 0.16 | 0.2581 | UNCHANGED |
| Dtk | 0.0035 | 1.00 | 1.56 | -0.56 | 0.12 | 0.0087 | UP |
| Eotaxin-1 | 0.1610 | 1.00 | 1.96 | -0.96 | 0.58 | 0.1699 | UNCHANGED |
| Eotaxin-2 | 0.0000 | 1.00 | 1.31 | -0.31 | 0.02 | 0.0001 | UP |
| E-Selectin | 0.0592 | 1.00 | 2.93 | -1.93 | 0.74 | 0.0854 | UNCHANGED |
| FAS ligand | 0.3612 | 1.00 | 1.70 | -0.70 | 0.68 | 0.3203 | UNCHANGED |
| Fc gamma RIIB | 0.1588 | 1.00 | 1.86 | -0.86 | 0.53 | 0.1699 | UNCHANGED |
| Flt3 Ligand | 0.0004 | 1.00 | 2.74 | -1.74 | 0.25 | 0.0015 | UP |
| Fractalkine | 0.3051 | 1.00 | 0.87 | 0.13 | 0.12 | 0.2838 | UNCHANGED |
| G-CSF | 0.5571 | 1.00 | 0.93 | 0.08 | 0.12 | 0.4580 | UNCHANGED |
| GITR | 0.0020 | 1.00 | 1.41 | -0.41 | 0.08 | 0.0057 | UP |
| GM-CSF | 0.0338 | 1.00 | 1.35 | -0.35 | 0.13 | 0.0530 | UP |
| Icam-1 | 0.0001 | 1.00 | 4.52 | -3.52 | 0.35 | 0.0003 | UP |
| IFN-gamma | 0.0000 | 1.00 | 2.73 | -1.73 | 0.08 | 0.0002 | UP |
| IGF-1 | 0.0003 | 1.00 | 3.20 | -2.20 | 0.30 | 0.0013 | UP |
| IGF-2 | 0.0016 | 1.00 | 1.88 | -0.88 | 0.16 | 0.0050 | UP |
| IGF-BP-2 | 0.1292 | 1.00 | 1.16 | -0.16 | 0.09 | 0.1522 | UNCHANGED |
| IGF-BP-3 | 0.0216 | 1.00 | 1.41 | -0.41 | 0.13 | 0.0363 | UP |
| IGF-BP-5 | 0.0036 | 1.00 | 1.66 | -0.66 | 0.14 | 0.0087 | UP |
| IGF-BP-6 | 0.1029 | 1.00 | 1.32 | -0.32 | 0.17 | 0.1322 | UNCHANGED |
| IL-10 | 0.3625 | 1.00 | 1.13 | -0.13 | 0.13 | 0.3203 | UNCHANGED |
| IL-12-p40/p70 | 0.0145 | 1.00 | 1.98 | -0.98 | 0.29 | 0.0262 | UP |
| IL-12-p70 | 0.3125 | 1.00 | 1.16 | -0.16 | 0.14 | 0.2869 | UNCHANGED |
| IL-13 | 0.1333 | 1.00 | 1.30 | -0.30 | 0.17 | 0.1545 | UNCHANGED |
| IL-15 | 0.0113 | 1.00 | 1.32 | -0.32 | 0.09 | 0.0228 | UP |

| | | | | | | | |
|---|---|---|---|---|---|---|---|
| IL-17 RB | 0.0992 | 1.00 | 1.63 | -0.63 | 0.32 | 0.1298 | UNCHANGED |
| IL-17A | 0.1462 | 1.00 | 1.16 | -0.16 | 0.09 | 0.1615 | UNCHANGED |
| IL1-alpha | 0.0000 | 1.00 | 1.57 | -0.57 | 0.04 | 0.0001 | UP |
| IL1-beta | 0.0000 | 1.00 | 2.47 | -1.47 | 0.08 | 0.0001 | UP |
| IL-2 | 0.0040 | 1.00 | 1.39 | -0.39 | 0.09 | 0.0092 | UP |
| IL-3 | 0.2856 | 1.00 | 1.24 | -0.24 | 0.20 | 0.2692 | UNCHANGED |
| IL-3 Rb | 0.0752 | 1.00 | 1.39 | -0.39 | 0.18 | 0.1042 | UNCHANGED |
| IL-4 | 0.0371 | 1.00 | 0.73 | 0.28 | 0.10 | 0.0562 | DOWN |
| IL-5 | 0.7307 | 1.00 | 1.06 | -0.06 | 0.17 | 0.5572 | UNCHANGED |
| IL-6 | 0.0167 | 1.00 | 1.80 | -0.79 | 0.24 | 0.0287 | UP |
| IL-7 | 0.0023 | 1.00 | 2.78 | -1.78 | 0.35 | 0.0063 | UP |
| IL-9 | 0.4428 | 1.00 | 0.90 | 0.10 | 0.12 | 0.3818 | UNCHANGED |
| iTAC | 0.0039 | 1.00 | 1.42 | -0.42 | 0.09 | 0.0092 | UP |
| KC | 0.0000 | 1.00 | 1.80 | -0.80 | 0.05 | 0.0001 | UP |
| Leptin | 0.5419 | 1.00 | 1.31 | -0.31 | 0.47 | 0.4561 | UNCHANGED |
| Leptin R | 0.4259 | 1.00 | 1.12 | -0.12 | 0.14 | 0.3717 | UNCHANGED |
| LIX | 0.3253 | 1.00 | 0.86 | 0.14 | 0.13 | 0.2949 | UNCHANGED |
| L-Selectin | 0.1127 | 1.00 | 1.34 | -0.34 | 0.18 | 0.1398 | UNCHANGED |
| Lungkine | 0.0000 | 1.00 | 1.74 | -0.74 | 0.07 | 0.0002 | UP |
| Lymphotactin | 0.0121 | 1.00 | 0.61 | 0.39 | 0.11 | 0.0230 | DOWN |
| MCP-1 | 0.9175 | 1.00 | 0.99 | 0.01 | 0.12 | 0.6757 | UNCHANGED |
| MCP-5 | 0.1408 | 1.00 | 0.81 | 0.19 | 0.11 | 0.1580 | UNCHANGED |
| M-CSF | 0.6673 | 1.00 | 0.95 | 0.06 | 0.12 | 0.5301 | UNCHANGED |
| MDC | 0.0067 | 1.00 | 3.62 | -2.62 | 0.65 | 0.0144 | UP |
| MIG | 0.1397 | 1.00 | 0.70 | 0.30 | 0.18 | 0.1580 | UNCHANGED |
| MIP-1-alpha | 0.0145 | 1.00 | 1.68 | -0.68 | 0.20 | 0.0262 | UP |
| MIP-1-gamma | 0.1068 | 1.00 | 1.50 | -0.50 | 0.26 | 0.1348 | UNCHANGED |
| MIP-2 | 0.0000 | 1.00 | 1.51 | -0.51 | 0.04 | 0.0001 | UP |
| MIP-3-alpha | 0.7610 | 1.00 | 1.14 | -0.15 | 0.46 | 0.5664 | UNCHANGED |
| MIP-3-beta | 0.0373 | 1.00 | 1.32 | -0.32 | 0.12 | 0.0562 | UP |
| MMP-2 | 0.0002 | 1.00 | 2.43 | -1.43 | 0.17 | 0.0009 | UP |
| MMP-3 | 0.0000 | 1.00 | 2.78 | -1.78 | 0.15 | 0.0002 | UP |
| OPG | 0.0008 | 1.00 | 1.54 | -0.53 | 0.09 | 0.0026 | UP |
| OPN | 0.0000 | 1.00 | 2.04 | -1.04 | 0.07 | 0.0001 | UP |
| PF4 | 0.0880 | 1.00 | 1.24 | -0.24 | 0.12 | 0.1173 | UNCHANGED |
| Pro-MMP-9 | 0.2039 | 1.00 | 0.06 | 0.94 | 0.31 | 0.2089 | UNCHANGED |
| P-Selectin | 0.0633 | 1.00 | 1.25 | -0.25 | 0.11 | 0.0894 | UNCHANGED |
| RANTES | 0.0004 | 1.00 | 1.47 | -0.48 | 0.07 | 0.0015 | UP |
| Resistin | 0.0518 | 1.00 | 1.65 | -0.64 | 0.27 | 0.0763 | UNCHANGED |
| SCF | 0.6805 | 1.00 | 0.86 | 0.15 | 0.34 | 0.5346 | UNCHANGED |

| | | | | | | | |
|---|---|---|---|---|---|---|---|
| SDF-1-alpha | 0.0092 | 1.00 | 0.52 | 0.48 | 0.13 | 0.0192 | DOWN |
| Shh-N | 0.0002 | 1.00 | 1.91 | -0.91 | 0.12 | 0.0010 | UP |
| sTNF RI | 0.0000 | 1.00 | 1.72 | -0.72 | 0.05 | 0.0001 | UP |
| sTNF RII | 0.0310 | 1.00 | 1.35 | -0.35 | 0.12 | 0.0498 | UP |
| TARC | 0.1201 | 1.00 | 0.82 | 0.18 | 0.10 | 0.1464 | UNCHANGED |
| TCK-1 | 0.5268 | 1.00 | 1.36 | -0.36 | 0.53 | 0.4488 | UNCHANGED |
| TECK | 0.0822 | 1.00 | 1.39 | -0.39 | 0.18 | 0.1118 | UNCHANGED |
| TIMP-1 | 0.0002 | 1.00 | 1.51 | -0.51 | 0.06 | 0.0010 | UP |
| TIMP-2 | 0.0000 | 1.00 | 2.30 | -1.30 | 0.11 | 0.0002 | UP |
| TNF-alpha | 0.0002 | 1.00 | 2.43 | -1.43 | 0.18 | 0.0009 | UP |
| TPO | 0.0119 | 1.00 | 1.64 | -0.64 | 0.18 | 0.0230 | UP |
| TRANCE | 0.1271 | 1.00 | 1.41 | -0.41 | 0.23 | 0.1522 | UNCHANGED |
| TROY | 0.0276 | 1.00 | 1.51 | -0.51 | 0.18 | 0.0453 | UP |
| TSLP | 0.0031 | 1.00 | 2.28 | -1.29 | 0.24 | 0.0082 | UP |
| VCAM-1 | 0.2385 | 1.00 | 1.13 | -0.13 | 0.10 | 0.2409 | UNCHANGED |
| VEGF R1 | 0.2853 | 1.00 | 2.26 | -1.26 | 1.03 | 0.2692 | UNCHANGED |
| VEGF R2 | 0.0018 | 1.00 | 3.84 | -2.84 | 0.53 | 0.0052 | UP |
| VEGF R3 | 0.0004 | 1.00 | 4.78 | -3.79 | 0.53 | 0.0015 | UP |
| VEGF-A | 0.7330 | 1.00 | 1.13 | -0.13 | 0.35 | 0.5572 | UNCHANGED |
| VEGF-D | 0.2746 | 1.00 | 1.38 | -0.38 | 0.26 | 0.2692 | UNCHANGED |

**Supplementary Table 1. Differentially secreted proteins in NTg astrocytes treated with PPIA compared to untreated NTg astrocytes.** List of alphabetically ordered 95 proteins analysed by cytokine array in media of untreated NTg astrocytes (N:U) and NTg astrocytes treated 24h with PPIA (N:P). Data are expressed as fold of N:U. Multiple unpaired T-Test. Difference is N:U – N:P. Significant differences were considered when $P < 0.05$. Table referrers to Figure 4.

#### Supplementary Table 2

| Pathways and relative proteins upregulated in N:P | | | |
|---|---|---|---|
| **Term** | **Nr. Proteins** | **Associated Proteins Found** | **Corresponding Factors** |
| Cytokine-cytokine receptor interaction | 26.00 | [Ccl19, Ccl22, Ccl24, Ccl3, Ccl5, Csf2, Cxcl1, Cxcl11, Cxcl15, Cxcl2, Ifng, Il12b, Il15, Il1a, Il1b, Il2, Il6, Il7, Thpo, Tnf, Tnfrsf11b, Tnfrsf18, Tnfrsf19, Tnfrsf1a, Tnfrsf1b, Tslp] | [MIP-3-beta, MDC, Eotaxin-2, MIP-1-alpha, RANTES, GM-CSF, KC, iTAC, Lungkine, MIP-2, IFN-gamma, IL12-p40/p70, IL-15, IL1-alpha, IL1-beta, IL-2, IL-6, IL-7, TPO, TNF-alpha, OPG, GITR, TROY, sTNF RI, sTNF RII, TSLP] |
| Signaling receptor binding | 22.00 | [Ccl19, Ccl24, Ccl3, Ccl5, Csf2, Cxcl15, Fgf2, Flt3l, Ifng, Igf1, Igf2, Il12b, Il15, Il1a, Il1b, Il2, Il6, Il7, Shh, Timp2, Tnf, Tslp] | [MIP-3-beta, Eotaxin-2, MIP-1-alpha, RANTES, GM-CSF, Lungkine, bFGF, FIt3 Ligand, IFN-gamma, IGF-1, IGF-2, IL12-p40/p70, IL-15, IL1-alpha, IL1-beta, IL-2, IL-6, IL-7, Shh-N, TIMP-2, TNF-alpha, TSLP] |
| Receptor ligand activity | 17.00 | [Ccl19, Ccl24, Ccl3, Csf2, Cxcl15, Fgf2, Ifng, Igf2, Il12b, Il15, Il1a, Il1b, Il2, Il6, Il7, Tnf, Tslp] | [MIP-3-beta, Eotaxin-2, MIP-1-alpha, GM-CSF, Lungkine, bFGF, IFN-gamma, IGF-2, IL12-p40/p70, IL-15, IL1-alpha, IL1-beta, IL-2, IL-6, IL-7, TNF-alpha, TSLP] |
| TNF signaling pathway | 12.00 | [Ccl5, Csf2, Cxcl1, Cxcl2, Icam1, Il15, Il1b, Il6, Mmp3, Tnf, Tnfrsf1a, Tnfrsf1b] | [RANTES, GM-CSF, KC, MIP-2, Icam-1, IL-15, IL1-beta, IL-6, MMP-3, TNF-alpha, sTNF RI, sTNF RII] |
| Leukocyte migration | 10.00 | [Ccl19, Ccl24, Ccl3, Ccl5, Cxcl15, Icam1, Ifng, Il1b, Spp1, Tnf] | [MIP-3-beta, Eotaxin-2, MIP-1-alpha, RANTES, Lungkine, Icam-1, IFN-gamma, IL1-beta, OPN, TNF-alpha] |
| Regulation of leukocyte differentiation | 10.00 | [Ccl19, Flt3l, Ifng, Il15, Il2, Il6, Il7, Shh, Tnf, Tnfrsf11b] | [MIP-3-beta, Flt3 Ligand, IFN-gamma, IL-15, IL-2, IL-6, IL-7, Shh-N, TNF-alpha, sTNF RII] |
| Positive regulation of lymphocyte activation | 9.00 | [Ccl19, Flt3l, Ifng, Il12b, Il15, Il2, Il6, Il7, Shh] | [MIP-3-beta, Flt3 Ligand, IFN-gamma, IL12-p40/p70, IL-15, IL-2, IL-6, IL-7, Shh-N] |
| Extrinsic apoptotic signaling pathway | 7.00 | [Igf1, Il1a, Il1b, Il2, Il7, Tnf, Tnfrsf1b] | [IGF-1, IL1-alpha, IL1-beta, IL-2, IL-7, TNF-alpha, sTNF RII] |
| NF-kappa B signaling pathway | 7.00 | [Ccl19, Cxcl1, Cxcl2, Icam1, Il1b, Tnf, Tnfrsf1a] | [MIP-3-beta, KC, MIP-2, Icam-1, IL1-beta, TNF-alpha, sTNF RI] |
| Regulation of adaptive immune response | 7.00 | [Ccl19, Ifng, Il12b, Il2, Il6, Tnf, Tnfrsf1b] | [MIP-3-beta, IFN-gamma, IL12-p40/p70, IL-2, IL-6, TNF-alpha, sTNF RII] |
| Regulation of lymphocyte differentiation | 7.00 | [Ccl19, Flt3l, Il15, Il2, Il6, Il7, Shh] | [MIP-3-beta, Flt3 Ligand, IL-15, IL-2, IL-6, IL-7, Shh-N] |
| Regulation of lymphocyte immunity | 6.00 | [Ifng, Il12b, Il2, Il6, Tnf, Tnfrsf1b] | [IFN-gamma, IL12-p40/p70, IL-2, IL-6, TNF-alpha, sTNF RII] |
| Regulation of protein kinase B signaling | 6.00 | [Fgf2, Igf1, Igfbp5, Mmp3, Thpo, Tnf] | [bFGF, IGF-1, IGF-BP-5, MMP-3, TPO, TNF-alpha] |
| Matrix metalloproteinases | 5.00 | [Mmp2, Mmp3, Timp1, Timp2, Tnf] | [MMP-2, MMP-3, TIMP-1, TIMP-2, TNF-alpha] |
| Positive regulation of angiogenesis | 5.00 | [Ccl24, Fgf2, Il1a, Il1b, Kdr] | [Eotaxin-2, bFGF, IL1-alpha, IL1-beta, VEGFR2] |
| Response to tumor necrosis factor | 5.00 | [Ccl5, Il6, Tnf, Tnfrsf1a, Tnfrsf1b] | [RANTES, IL-6, TNF-alpha, sTNF RI, sTNF RII] |

**Supplementary Table 2. Pathways and relative proteins upregulated in NTg astrocytes treated with PPIA.** List of pathways (terms) and proteins, related to the single pathway, found by

ClueGO analysis of the 43 upregulated proteins in NTg astrocytes treated with PPIA. Table referrers to Figure 4.

**Supplementary Table 3**

| NTg untreated (N:U) vs SOD1G93A untreated (G:U) | | | | | | | |
|---|---|---|---|---|---|---|---|
| | | Mean | | | SE of | | |
| **Factors** | **P value** | **N:U** | **G:U** | **Difference** | **difference** | **q value** | **in G:U** |
| axl | 0.0671 | 1.00 | 1.43 | -0.43 | 0.19 | 0.0511 | UNCHANGED |
| bFGF | 0.0001 | 1.00 | 2.41 | -1.41 | 0.15 | 0.0002 | UP |
| BLC | 0.0635 | 1.00 | 1.37 | -0.37 | 0.15 | 0.0491 | UNCHANGED |
| CCL1 | 0.0061 | 1.00 | 1.43 | -0.43 | 0.10 | 0.0062 | UP |
| CD26 | 0.0204 | 1.00 | 3.68 | -2.67 | 0.80 | 0.0171 | UP |
| CD30/TNFRSF8 | 0.0883 | 1.00 | 1.36 | -0.36 | 0.18 | 0.0652 | UNCHANGED |
| CD30L | 0.2389 | 1.00 | 1.14 | -0.14 | 0.11 | 0.1535 | UNCHANGED |
| CD40 | 0.4694 | 1.00 | 1.14 | -0.13 | 0.17 | 0.2640 | UNCHANGED |
| CRG-2 | 0.0140 | 1.00 | 1.40 | -0.40 | 0.12 | 0.0128 | UP |
| CTACK | 0.0013 | 1.00 | 2.13 | -1.13 | 0.20 | 0.0018 | UP |
| CXCL16 | 0.0040 | 1.00 | 1.61 | -0.61 | 0.13 | 0.0044 | UP |
| Dtk | 0.0000 | 1.00 | 2.46 | -1.46 | 0.07 | 0.0000 | UP |
| Eotaxin-1 | 0.1895 | 1.00 | 1.65 | -0.64 | 0.43 | 0.1302 | UNCHANGED |
| Eotaxin-2 | 0.0000 | 1.00 | 1.56 | -0.56 | 0.04 | 0.0000 | UP |
| E-Selectin | | 1.00 | | | | | not detected |
| FAS ligand | 0.9825 | 1.00 | 1.02 | -0.02 | 0.84 | 0.5173 | UNCHANGED |
| Fc gamma RIIB | 0.0034 | 1.00 | 4.77 | -3.78 | 0.81 | 0.0039 | UP |
| Flt3 Ligand | 0.0000 | 1.00 | 4.43 | -3.43 | 0.28 | 0.0000 | UP |
| Fractalkine | 0.4431 | 1.00 | 1.08 | -0.08 | 0.09 | 0.2520 | UNCHANGED |
| G-CSF | 0.8615 | 1.00 | 0.97 | 0.03 | 0.15 | 0.4634 | UNCHANGED |
| GITR | 0.0000 | 1.00 | 1.69 | -0.69 | 0.06 | 0.0001 | UP |
| GM-CSF | 0.0016 | 1.00 | 1.55 | -0.55 | 0.10 | 0.0020 | UP |
| Icam-1 | 0.0000 | 1.00 | 6.70 | -5.70 | 0.36 | 0.0000 | UP |
| IFN-gamma | <0.000001 | 1.00 | 3.78 | -2.78 | 0.08 | 0.0000 | UP |
| IGF-1 | 0.0000 | 1.00 | 5.58 | -4.58 | 0.28 | 0.0000 | UP |
| IGF-2 | 0.0020 | 1.00 | 1.98 | -0.98 | 0.19 | 0.0024 | UP |
| IGF-BP-2 | 0.0000 | 1.00 | 2.09 | -1.09 | 0.10 | 0.0001 | UP |
| IGF-BP-3 | 0.0013 | 1.00 | 1.57 | -0.57 | 0.10 | 0.0018 | UP |
| IGF-BP-5 | 0.0003 | 1.00 | 1.80 | -0.80 | 0.10 | 0.0005 | UP |
| IGF-BP-6 | 0.0086 | 1.00 | 1.53 | -0.53 | 0.14 | 0.0085 | UP |
| IL-10 | 0.3656 | 1.00 | 1.24 | -0.24 | 0.25 | 0.2154 | UNCHANGED |
| IL-12-p40/p70 | 0.0053 | 1.00 | 1.85 | -0.85 | 0.20 | 0.0057 | UP |
| IL-12-p70 | 0.0210 | 1.00 | 1.33 | -0.33 | 0.11 | 0.0171 | UP |
| IL-13 | 0.1805 | 1.00 | 1.19 | -0.19 | 0.13 | 0.1276 | UNCHANGED |
| IL-15 | 0.0020 | 1.00 | 5.78 | -4.78 | 0.92 | 0.0024 | UP |
| IL-17 RB | 0.0747 | 1.00 | 4.89 | -3.89 | 1.62 | 0.0560 | UNCHANGED |

| | | | | | | | |
|---|---|---|---|---|---|---|---|
| IL-17A | 0.0013 | 1.00 | 1.43 | -0.43 | 0.08 | 0.0018 | UP |
| IL-1-alpha | <0.000001 | 1.00 | 1.84 | -0.84 | 0.03 | 0.0000 | UP |
| IL-1-beta | 0.0009 | 1.00 | 3.08 | -2.08 | 0.34 | 0.0014 | UP |
| IL-2 | 0.0211 | 1.00 | 1.45 | -0.45 | 0.14 | 0.0171 | UP |
| IL-3 | 0.1856 | 1.00 | 1.49 | -0.49 | 0.32 | 0.1293 | UNCHANGED |
| IL-3 Rb | 0.0007 | 1.00 | 2.15 | -1.15 | 0.18 | 0.0012 | UP |
| IL-4 | 0.8582 | 1.00 | 1.02 | -0.02 | 0.09 | 0.4634 | UNCHANGED |
| IL-5 | 0.3878 | 1.00 | 0.80 | 0.21 | 0.22 | 0.2255 | UNCHANGED |
| IL-6 | 0.0081 | 1.00 | 2.00 | -0.99 | 0.26 | 0.0081 | UP |
| IL-7 | 0.0145 | 1.00 | 4.20 | -3.20 | 0.94 | 0.0131 | UP |
| IL-9 | 0.2032 | 1.00 | 1.15 | -0.15 | 0.11 | 0.1348 | UNCHANGED |
| iTAC | <0.000001 | 1.00 | 2.94 | -1.94 | 0.07 | 0.0000 | UP |
| KC | <0.000001 | 1.00 | 2.22 | -1.22 | 0.04 | 0.0000 | UP |
| Leptin | 0.2721 | 1.00 | 1.63 | -0.63 | 0.49 | 0.1683 | UNCHANGED |
| Leptin R | 0.0134 | 1.00 | 1.51 | -0.51 | 0.15 | 0.0125 | UP |
| LIX | 0.0105 | 1.00 | 1.42 | -0.42 | 0.11 | 0.0101 | UP |
| L-Selectin | 0.0002 | 1.00 | 2.49 | -1.49 | 0.18 | 0.0003 | UP |
| Lungkine | <0.000001 | 1.00 | 3.03 | -2.03 | 0.05 | 0.0000 | UP |
| Lymphotactin | 0.5173 | 1.00 | 0.93 | 0.08 | 0.11 | 0.2876 | UNCHANGED |
| MCP-1 | 0.2043 | 1.00 | 1.14 | -0.14 | 0.10 | 0.1348 | UNCHANGED |
| MCP-5 | 0.0344 | 1.00 | 0.75 | 0.25 | 0.09 | 0.0270 | DOWN |
| M-CSF | 0.2712 | 1.00 | 1.12 | -0.12 | 0.09 | 0.1683 | UNCHANGED |
| MDC | 0.0000 | 1.00 | 5.14 | -4.14 | 0.31 | 0.0000 | UP |
| MIG | 0.0198 | 1.00 | 1.64 | -0.64 | 0.17 | 0.0169 | UP |
| MIP-1-alpha | 0.0126 | 1.00 | 2.90 | -1.90 | 0.54 | 0.0120 | UP |
| MIP-1-gamma | 0.1248 | 1.00 | 1.64 | -0.63 | 0.35 | 0.0908 | UNCHANGED |
| MIP-2 | 0.0000 | 1.00 | 1.61 | -0.61 | 0.04 | 0.0000 | UP |
| MIP-3-alpha | 0.3918 | 1.00 | 1.39 | -0.40 | 0.43 | 0.2255 | UNCHANGED |
| MIP-3-beta | 0.0015 | 1.00 | 1.54 | -0.54 | 0.10 | 0.0019 | UP |
| MMP-2 | 0.0000 | 1.00 | 3.11 | -2.11 | 0.15 | 0.0000 | UP |
| MMP-3 | <0.000001 | 1.00 | 4.93 | -3.93 | 0.18 | 0.0000 | UP |
| OPG | 0.0001 | 1.00 | 1.92 | -0.92 | 0.09 | 0.0001 | UP |
| OPN | <0.000001 | 1.00 | 2.02 | -1.02 | 0.03 | 0.0000 | UP |
| PF4 | 0.0015 | 1.00 | 1.63 | -0.63 | 0.11 | 0.0019 | UP |
| Pro-MMP-9 | 0.2191 | 1.00 | 1.87 | -0.87 | 0.31 | 0.1426 | UNCHANGED |
| P-Selectin | 0.0002 | 1.00 | 1.74 | -0.75 | 0.09 | 0.0003 | UP |
| RANTES | 0.0000 | 1.00 | 2.05 | -1.05 | 0.08 | 0.0000 | UP |
| Resistin | 0.0009 | 1.00 | 5.09 | -4.08 | 0.67 | 0.0014 | UP |
| SCF | 0.2870 | 1.00 | 1.57 | -0.56 | 0.48 | 0.1753 | UNCHANGED |
| SDF-1-alpha | 0.3415 | 1.00 | 1.18 | -0.18 | 0.18 | 0.2061 | UNCHANGED |

| | | | | | | | |
|---|---|---|---|---|---|---|---|
| Shh-N | 0.6250 | 1.00 | 1.13 | -0.13 | 0.25 | 0.3437 | UNCHANGED |
| sTNF RI | 0.0000 | 1.00 | 1.97 | -0.98 | 0.07 | 0.0000 | UP |
| sTNF RII | 0.0054 | 1.00 | 1.45 | -0.46 | 0.11 | 0.0057 | UP |
| TARC | 0.8837 | 1.00 | 0.99 | 0.02 | 0.10 | 0.4703 | UNCHANGED |
| TCK-1 | 0.3459 | 1.00 | 0.59 | 0.41 | 0.39 | 0.2062 | UNCHANGED |
| TECK | 0.0195 | 1.00 | 1.54 | -0.54 | 0.17 | 0.0169 | UP |
| TIMP-1 | 0.0013 | 1.00 | 1.37 | -0.37 | 0.07 | 0.0018 | UP |
| TIMP-2 | 0.0000 | 1.00 | 3.77 | -2.77 | 0.26 | 0.0001 | UP |
| TNF-alpha | 0.0000 | 1.00 | 3.59 | -2.59 | 0.19 | 0.0000 | UP |
| TPO | 0.0008 | 1.00 | 1.87 | -0.87 | 0.14 | 0.0014 | UP |
| TRANCE | 0.0309 | 1.00 | 2.69 | -1.69 | 0.60 | 0.0247 | UP |
| TROY | 0.0001 | 1.00 | 3.16 | -2.16 | 0.25 | 0.0003 | UP |
| TSLP | 0.0019 | 1.00 | 2.78 | -1.78 | 0.30 | 0.0023 | UP |
| VCAM-1 | 0.0033 | 1.00 | 1.47 | -0.47 | 0.10 | 0.0038 | UP |
| VEGF R1 | 0.2612 | 1.00 | 2.49 | -1.49 | 1.18 | 0.1657 | UNCHANGED |
| VEGF R2 | 0.0195 | 1.00 | 3.48 | -2.48 | 0.78 | 0.0169 | UP |
| VEGF R3 | 0.0013 | 1.00 | 5.37 | -4.37 | 0.77 | 0.0018 | UP |
| VEGF-A | 0.2007 | 1.00 | 1.38 | -0.38 | 0.26 | 0.1348 | UNCHANGED |
| VEGF-D | 0.1328 | 1.00 | 0.10 | 0.90 | 0.19 | 0.0953 | UNCHANGED |

**Supplementary Table 3. Differentially secreted proteins in untreated SOD1$^{G93A}$ astrocytes compared to untreated NTg astrocytes.** List of alphabetically ordered 95 proteins analysed by cytokine array in media of untreated NTg (N:U) and SOD1$^{G93A}$ astrocytes (G:U). Data are expressed as fold of N:U. Multiple unpaired T-Test. Difference is N:U – G:U. Significant differences were considered when $P < 0.05$. Table referrers to Figure 5.

**Supplementary Table 4**

| Upregulated proteins | | |
|---|---|---|
| **in N:P only** | **in N:P and G:U** | **in G:U only** |
| Shh-N | bFGF | IGF BP-2 |
| | CD26 | CCL1 |
| | Dtk | CRG-2 |
| | Eotaxin-2 | CTACK |
| | Flt3 Ligand | CXCL16 |
| | GITR | Fc gamma RIIB |
| | GM-CSF | IGF-BP-6 |
| | Icam-1 | IL-12-p70 |
| | IFN-gamma | IL-17A |
| | IGF-1 | IL-3 Rb |
| | IGF-2 | Leptin R |
| | IGF-BP-3 | LIX |
| | IGF-BP-5 | L-Selectin |
| | IL-12-p40/p70 | MIG |
| | IL-15 | PF4 |
| | IL1-alpha | P-Selectin |
| | IL1-beta | Resistin |
| | IL-2 | TECK |
| | IL-6 | TRANCE |
| | IL-7 | VCAM-1 |
| | iTAC | |
| | KC | |
| | Lungkine | |
| | MDC | |
| | MIP-1-alpha | |
| | MIP-2 | |
| | MIP-3-beta | |
| | MMP-2 | |
| | MMP-3 | |
| | OPG | |
| | OPN | |
| | RANTES | |
| | sTNF RI | |
| | sTNF RII | |
| | TIMP-1 | |
| | TIMP-2 | |
| | TNF-alpha | |

| | |
|---|---|
| | TPO |
| | TROY |
| | TSLP |
| | VEGFR2 |
| | VEGFR3 |

**Supplementary Table 4. Commonly secreted proteins in SOD1$^{G93A}$ astrocytes and NTg astrocytes treated with PPIA.** List of alphabetically ordered proteins upregulated in NTg astrocytes treated with PPIA (N:P) only, in untreated SOD1$^{G93A}$ astrocytes (G:U) only, or commonly upregulated in both conditions. Table referrers to Figure 5.

### Supplementary Table 5

| Pathways and relative proteins upregulated in G:U | | | |
|---|---|---|---|
| **Term** | **Nr. Proteins** | **Associated Proteins Found** | **Corresponding Factors** |
| Cytokine-cytokine receptor interaction | 39.00 | [Ccl1, Ccl19, Ccl22, Ccl24, Ccl25, Ccl27a, Ccl3, Ccl5, Csf2, Csf2rb, Cxcl1, Cxcl10, Cxcl11, Cxcl15, Cxcl16, Cxcl2, Cxcl5, Cxcl9, Ifng, Il12a, Il12b, Il15, Il17a, Il1a, Il1b, Il2, Il6, Il7, Lepr, Pf4, Thpo, Tnf, Tnfrsf11b, Tnfrsf18, Tnfrsf19, Tnfrsf1a, Tnfrsf1b, Tnfsf11, Tslp] | [CCL1, MIP-3-beta,MDC,Eotaxin-2,TECK, CTACK, MIP-1-alpha, RANTES, GM-CSF, IL3 Rb, KC, CRG-2, iTAC, Lungkine, CXCL16, MIP-2, LIX, MIG, IFN-gamma, IL12-p70, IL12-p40/p70, IL-15, IL-17A, IL1-alpha, IL1-beta, IL-2, IL-6, IL-7, Leptin R, PF4, TPO, TNF-alpha, OPG, GITR, TROY, sTNF RI, sTNF RII, TRANCE, TSLP] |
| Signaling receptor binding | 27.00 | [Ccl1, Ccl19, Ccl24, Ccl25, Ccl27a, Ccl3, Ccl5, Csf2, Cxcl15, Cxcl16, Fgf2, Flt3l, Ifng, Igf1, Igf2, Il12a, Il12b, Il15, Il1a, Il1b, Il2, Il6, Il7, Timp2, Tnf, Tnfsf11, Tslp] | [CCL1, MIP-3-beta, Eotaxin-2, TECK, CTACK, MIP-1-alpha, RANTES, GM-CSF, Lungkine, CXCL16, bFGF, Flt3 Ligand, IFN-gamma, IGF-2, IL12-p70, IL12-p40/p70, IL-15, IL1-alpha, IL1-beta, IL-2, IL-6, IL-7, TIMP-2, TNF-alpha, TRANCE, TSLP] |
| Receptor ligand activity | 22.00 | [Ccl1, Ccl19, Ccl24, Ccl25, Ccl27a, Ccl3, Csf2, Cxcl15, Cxcl16, Fgf2, Ifng, Igf2, Il12a, Il12b, Il15, Il1a, Il1b, Il2, Il6, Il7, Tnf, Tslp] | [CCL1, MIP-3-beta, Eotaxin-2, TECK, CTACK, MIP-1-alpha, GM-CSF, Lungkine, CXCL16, bFGF, IFN-gamma, IGF-2, IL12-p70, IL12-p40/p70, IL-15, IL1-alpha, IL1-beta, IL-2, IL-6, IL-7, TNF-alpha, TSLP] |
| Regulation of defense response | 16.00 | [Ccl24, Ccl3, Cxcl1, Cxcl5, Fcgr2b, Ifng, Il12b, Il15, Il17a, Il1b, Il2, Tnf, Tnfrsf1a, Tnfrsf1b, Tnfsf11, Tyro3] | [Eotaxin-2, MIP-1-alpha, KC, Lungkine, Fc gamma RIIB, IFN-gamma, IL12-p40/p70, IL-15, IL-17A, IL1-beta, IL-2, TNF-alpha, sTNF RI, sTNF RII, TRANCE, Dtk] |
| TNF signaling pathway | 15.00 | [Ccl5, Csf2, Cxcl1, Cxcl10, Cxcl2, Cxcl5, Icam1, Il15, Il1b, Il6, Mmp3, Tnf, Tnfrsf1a, Tnfrsf1b, Vcam1] | [RANTES, GM-CSF, KC, CRG-2, MIP-2, LIX, ICAM-1, IL-15, IL1-beta, IL-6, MMP-3, TNF-alpha, sTNF RI, sTNF RII, VCAM-1] |
| Lipid and atherosclerosis | 13.00 | [Ccl3, Ccl5, Cxcl1, Cxcl2, Icam1, Il12a, Il12b, Il1b, Il6, Mmp3, Tnf, Tnfrsf1a, Vcam1] | [MIP-1-alpha, RANTES, KC, ICAM-1, IL12-p70, IL12-p40/p70, IL1-beta, IL-6, MMP-3, TNF-alpha, sTNF RI, VCAM-1] |
| Cytokines and Inflammatory response | 12.00 | [Csf2, Cxcl1, Ifng, Il12a, Il12b, Il15, Il1a, Il1b, Il2, Il6, Il7, Tnf] | [GM-CSF, KC, IFN-gamma, IL12-p70, IL12-p40/p70, IL-15, IL1-alpha, IL1-beta, IL-2, IL-6, IL-7, TNF-alpha] |
| Regulation of inflammatory response | 11.00 | [Ccl24, Ccl3, Fcgr2b, Il12b, Il1b, Il2, Tnf, Tnfrsf1a, Tnfrsf1b, Tnfsf11, Tyro3] | [Eotaxin-2, MIP-1-alpha, Fc gamma RIIB, Il12-p40/p70, IL1-beta, IL-2, TNF-alpha, sTNF RI, sTNF RII, TRANCE, Dtk] |
| IL-17 signaling pathway | 11.00 | [Csf2, Cxcl1, Cxcl10, Cxcl2, Cxcl5, Ifng, Il17a, Il1b, Il6, Mmp3, Tnf] | [GM-CSF, KC, CRG-2, MIP-2, LIX, IFN-gamma, IL-17A, IL1-beta, IL-6, MMP-3, TNF-alpha] |
| Regulation of response to external stimulus | 10.00 | [Ccl19, Ccl24, Ccl3, Ccl5, Cxcl1, Il17a, Il1b, Tnf, Tnfrsf1a, Tnfsf11] | [MIP-3-beta, Eotaxin-2, MIP-1-alpha, RANTES, KC, IL-17A, IL1-beta, TNF-alpha, sTNF RI, TRANCE] |
| Cytokine-mediated signaling pathway | 9.00 | [Csf2rb, Ifng, Il6, Lepr, Thpo, Tnf, Tnfrsf1a, Tnfrsf1b, Tnfsf11] | [IL3 RB, IFN-gamma, IL-6, Leptin R, TPO, TNF-alpha, sTNF RI, sTNF RII, TRANCE] |
| Regulation of leukocyte immunity | 9.00 | [Cxcl1, Cxcl5, Fcgr2b, Ifng, Il12b, Il2, Il6, Tnf, Tnfrsf1b] | [KC, LIX, FC gamma RIIB, IFN-gamma, IL12-p40/p70, IL-2, IL-6, TNF-alpha, sTNF RII] |
| Myeloid leukocyte migration | 8.00 | [Ccl24, Ccl3, Ccl5, Cxcl15, Ifng, Il17a, Il1b, Spp1] | [Eotaxin-2, MIP-1-alpha, RANTES, Lungkine, IFN-gamma, IL-17A, IL1-beta, OPN] |
| Positive regulation of immune effector | 8.00 | [Ccl19, Cxcl1, Ifng, Il12b, Il17a, Il2, Il6, Tnf] | [MIP-3-beta, KC, IFN-gamma, IL12-p40/p70, IL-17A, IL-2, IL-6, TNF-alpha] |
| Positive regulation of leukocyte proliferation | 8.00 | [Ccl19, Flt3l, Ifng, Il12a, Il12b, Il15, Il2, Il7] | [MIP-3-beta, Flt3 Ligand, IFN-gamma, IL12-p70, IL12-p40/p70, IL-15, IL-2, IL-7] |
| Regulation of adaptive immune response | 8.00 | [Ccl19, Fcgr2b, Ifng, Il12b, Il2, Il6, Tnf, Tnfrsf1b] | [MIP-3-beta, Fc gamma RIIB, IFN-gamma, IL12-p40/p70, IL-2, IL-6, TNF-alpha, sTNF RII] |
| Regulation of response to biotic stimulus | 8.00 | [Cxcl1, Cxcl5, Ifng, Il12b, Il15, Il17a, Il1b, Tyro3] | [KC, LIX, IFN-gamma, IL12-p40/p70, IL-15, IL-17A, IL1-beta, Dtk] |
| Extrinsic apoptotic signaling pathway | 7.00 | [Igf1, Il1a, Il1b, Il2, Il7, Tnf, Tnfrsf1b] | [IFG-1, IL1-alpha, IL1-beta, IL-2, IL-7, TNF-alpha, sTNF RII] |
| Regulation of lymphocyte immunity | 7.00 | [Fcgr2b, Ifng, Il12b, Il2, Il6, Tnf, Tnfrsf1b] | [Fc gamma RIIB, IFN-gamma, IL12-p40/p70, IL-2, IL-6, TNF-alpha, sTNF RII] |
| Regulation of mediator of immune response | 7.00 | [Fcgr2b, Ifng, Il17a, Il2, Il6, Tnf, Tnfrsf1b] | [Fc gamma RIIB, IFN-gamma, IL-17A, IL-2, IL-6, TNF-alpha, sTNF RII] |
| Regulation of protein kinase B signaling | 7.00 | [Fgf2, Igf1, Igfbp5, Mmp3, Thpo, Tnf, Tnfsf11] | [bFGF, IGF-1, IGF-BP-5, MMP-3, TPO, TNF-alpha, TRANCE] |
| Regulation of T cell proliferation | 7.00 | [Ccl19, Ifng, Il12a, Il12b, Il15, Il2, Tnfrsf1b] | [MIP-3-beta, IFN-gamma, IL12-p70, IL12-p40/p70, IL-15, IL-2, sTNF RII] |
| Response to virus | 7.00 | [Cxcl10, Cxcl9, Ifng, Il12b, Il15, Il1b, Il6] | [CRG-2, MIG, IFN-gamma, IL12-p40/p70, IL-15, IL1-beta, IL-16] |
| Insulin-like growth factor (IGF1)-Akt signaling | 7.00 | [Igf1, Igfbp2, Igfbp3, Igfbp5, Igfbp6, Tnf, Tnfrsf1a] | [IGF-1, IGF-BP-2, IGF-BP-3, IGF-BP-5, IFGBP-6, TNF-alpha, sTNF RI] |
| Response to tumor necrosis factor | 6.00 | [Ccl5, Il6, Tnf, Tnfrsf1a, Tnfrsf1b, Tnfsf11] | [RANTES, IL-6, TNF-alpha, sTNF RI, sTNF RII, TRANCE] |
| Positive regulation of angiogenesis | 5.00 | [Ccl24, Fgf2, Il1a, Il1b, Kdr] | [Eotaxin-2, bFGF, IL1-alpha, IL1-beta, VEGFR2] |
| Matrix metalloproteinases | 5.00 | [Mmp2, Mmp3, Timp1, Timp2, Tnf] | [MMP-2, MMP-3, TIMP-1, TIMP-2, TNF-alpha] |
| Type II interferon signaling (IFNG) | 5.00 | [Cxcl10, Cxcl9, Icam1, Ifng, Il1b] | [CRG-2, MIG, ICAM-1, IFN-gamma, IL1-beta] |

**Supplementary Table 5. Pathways and relative proteins upregulated in SOD1$^{G93A}$ astrocytes.** List of pathways (terms) and proteins, related to the single pathway, found by ClueGO analysis of the 62 upregulated proteins in untreated SOD1$^{G93A}$ astrocytes. Table referrers to Figure 5.

**Supplementary Table 6**

| SOD1G93A + CTR Ab (G:C) vs SOD1G93A + EMN Ab (G:E) | | | | | | | |
|---|---|---|---|---|---|---|---|
| | | Mean | | | SE of | | |
| Factors | P value | G:C | G:U | Difference | difference | q value | in G:E |
| bFGF | 0.0041 | 2.16 | 1.22 | 0.95 | 0.21 | 0.0037 | DOWN |
| CD26 | 0.0000 | 2.45 | 0.76 | 1.69 | 0.10 | 0.0000 | DOWN |
| Dtk | 0.0135 | 2.32 | 1.56 | 0.76 | 0.22 | 0.0109 | DOWN |
| Eotaxin-2 | <0.000001 | 1.79 | 0.60 | 1.19 | 0.04 | 0.0000 | DOWN |
| Flt3 Ligand | 0.0141 | 3.72 | 1.95 | 1.77 | 0.52 | 0.0110 | DOWN |
| GITR | 0.0000 | 1.74 | 0.59 | 1.15 | 0.11 | 0.0001 | DOWN |
| GM-CSF | 0.1219 | 1.63 | 1.26 | 0.38 | 0.21 | 0.0724 | UNCHANGED |
| Icam-1 | 0.0062 | 4.65 | 1.66 | 2.99 | 0.72 | 0.0055 | DOWN |
| IFN-gamma | 0.1991 | 3.00 | 2.04 | 0.96 | 0.65 | 0.1087 | UNCHANGED |
| IGF-1 | 0.0000 | 5.18 | 1.62 | 3.56 | 0.27 | 0.0001 | DOWN |
| IGF-2 | 0.0002 | 2.34 | 1.02 | 1.32 | 0.16 | 0.0003 | DOWN |
| IGF-BP-3 | 0.0205 | 1.55 | 0.74 | 0.81 | 0.26 | 0.0147 | DOWN |
| IGF-BP-5 | 0.0305 | 2.24 | 1.47 | 0.77 | 0.27 | 0.0199 | DOWN |
| IL-12-p40/p70 | 0.0239 | 2.46 | 1.39 | 1.06 | 0.35 | 0.0161 | DOWN |
| IL-15 | 0.0621 | 2.45 | 1.26 | 1.19 | 0.52 | 0.0392 | UNCHANGED |
| IL-1-alpha | 0.0002 | 1.75 | 1.28 | 0.47 | 0.06 | 0.0003 | DOWN |
| IL-1-beta | 0.0034 | 3.72 | 2.23 | 1.50 | 0.32 | 0.0033 | DOWN |
| IL-2 | 0.8100 | 1.33 | 1.40 | -0.07 | 0.26 | 0.3896 | UNCHANGED |
| IL-6 | 0.0001 | 2.05 | 1.24 | 0.81 | 0.09 | 0.0003 | DOWN |
| IL-7 | 0.0211 | 1.33 | 5.55 | -4.22 | 1.36 | 0.0147 | UP |
| iTAC | 0.0128 | 1.73 | 1.40 | 0.34 | 0.10 | 0.0108 | DOWN |
| KC | 0.0011 | 2.22 | 1.59 | 0.63 | 0.11 | 0.0012 | DOWN |
| Lungkine | 0.0022 | 1.56 | 1.11 | 0.46 | 0.09 | 0.0024 | DOWN |
| MDC | 0.0000 | 4.28 | 0.99 | 3.29 | 0.30 | 0.0001 | DOWN |
| MIP-1-alpha | 0.6227 | 2.20 | 2.39 | -0.19 | 0.36 | 0.3068 | UNCHANGED |
| MIP-2 | 0.0004 | 1.58 | 1.35 | 0.23 | 0.03 | 0.0005 | DOWN |
| MIP-3-beta | 0.0943 | 1.45 | 1.08 | 0.37 | 0.19 | 0.0577 | UNCHANGED |
| MMP-2 | 0.0032 | 1.85 | 0.90 | 0.95 | 0.20 | 0.0032 | DOWN |
| MMP-3 | 0.0003 | 3.57 | 1.13 | 2.44 | 0.32 | 0.0004 | DOWN |
| OPG | 0.0002 | 1.62 | 0.91 | 0.72 | 0.09 | 0.0003 | DOWN |
| OPN | 0.0001 | 2.05 | 0.80 | 1.25 | 0.13 | 0.0001 | DOWN |
| RANTES | 0.0000 | 2.09 | 0.56 | 1.53 | 0.12 | 0.0001 | DOWN |
| sTNF RI | <0.000001 | 2.39 | 1.45 | 0.95 | 0.04 | 0.0000 | DOWN |
| sTNF RII | 0.2281 | 1.28 | 1.02 | 0.26 | 0.20 | 0.1213 | UNCHANGED |
| TIMP-1 | 0.0004 | 1.70 | 0.92 | 0.78 | 0.11 | 0.0005 | DOWN |
| TIMP-2 | 0.5057 | 2.06 | 2.19 | -0.12 | 0.17 | 0.2554 | UNCHANGED |

| | | | | | | | |
|---|---|---|---|---|---|---|---|
| TNF-alpha | 0.0000 | 3.95 | 2.69 | 1.26 | 0.12 | 0.0001 | DOWN |
| TPO | 0.0190 | 1.86 | 1.14 | 0.72 | 0.23 | 0.0142 | DOWN |
| TROY | 0.1444 | 1.00 | 1.33 | -0.33 | 0.20 | 0.0833 | UNCHANGED |
| TSLP | 0.3612 | 2.48 | 1.76 | 0.73 | 0.73 | 0.1871 | UNCHANGED |
| VEGF R2 | 0.0002 | 3.48 | 0.23 | 3.25 | 0.35 | 0.0004 | DOWN |
| VEGF R3 | 0.1751 | 1.31 | 0.63 | 0.67 | 0.38 | 0.0983 | UNCHANGED |

**Supplementary Table 6. Differentially secreted proteins in SOD1$^{G93A}$ astrocytes treated with anti-EMMPRIN antibody compared to SOD1$^{G93A}$ astrocytes treated with a control antibody.** List of alphabetically ordered 42 proteins analysed by cytokine array in media of SOD1$^{G93A}$ astrocytes treated 24h with control antibody (G:C) or anti-EMMPIN antibody (G:E). Data are expressed as fold of N:U. Multiple unpaired T-Test. Difference is G:C – G:E. Significant differences were considered when $P < 0.05$. Table referrers to Figure 6.

#### Supplementary Table 7

| Pathways and relative proteins downregulated in G:E | | | |
|---|---|---|---|
| **Term** | **Nr. Proteins** | **Associated Proteins Found** | **Corresponding Factors** |
| Cytokine-cytokine receptor interaction | 17.00 | [Ccl22, Ccl24, Ccl5, Cxcl1, Cxcl11, Cxcl15, Cxcl2, Il12b, Il1a, Il1b, Il6, Thpo, Tnf, Tnfrsf11b, Tnfrsf18, Tnfrsf1a, Tnfrsf1b] | [MDC, Eotaxin-2, RANTES, KC, iTAC, Lungkine, MIP-2, IL12-p40/p70, IL1-alpha, IL1-beta, IL-6, TPO, TNF-alpha, OPG, GITR, sTNF RI, sTNF RII] |
| Lipid and atherosclerosis | 10.00 | [Ccl5, Cxcl1, Cxcl2, Icam1, Il12b, Il1b, Il6, Mmp3, Tnf, Tnfrsf1a] | [RANTES, KC, MIP-2, ICAM-1, IL12-p40/p70, IL1-beta, IL-6, MMP-3, TNF-alpha, sTNF RI] |
| TNF signaling pathway | 10.00 | [Ccl5, Cxcl1, Cxcl2, Icam1, Il1b, Il6, Mmp3, Tnf, Tnfrsf1a, Tnfrsf1b] | [RANTES, KC, MIP-2, ICAM-1, IL1-beta, IL-6, MMP-3, TNF-alpha, sTNF RI, sTNF RII] |
| Regulation of protein kinase B signaling | 6.00 | [Fgf2, Igf1, Igfbp5, Mmp3, Thpo, Tnf] | [bFGF, IGF-1, IGF-BP-5, MMP-3, TPO, TNF-alpha] |
| Myeloid leukocyte migration | 5.00 | [Ccl24, Ccl5, Cxcl15, Il1b, Spp1] | [Eotaxin-2, RANTES, Lungkine, IL1-beta, OPN] |
| Positive regulation of angiogenesis | 5.00 | [Ccl24, Fgf2, Il1a, Il1b, Kdr] | [Eotaxin-2, bFGF, IL1-alpha, IL1-beta, VEGFR2] |
| Response to tumor necrosis factor | 5.00 | [Ccl5, Il6, Tnf, Tnfrsf1a, Tnfrsf1b] | [RANTES, IL-6, TNF-alpha, sTNF RI, sTNF RII] |

**Supplementary Table 7. Pathways and relative proteins downregulated in SOD1$^{G93A}$ astrocytes after anti-EMMPRIN antibody treatment.** List of pathways (terms) and proteins, related to the single pathway, found by ClueGO analysis of the 42 downregulated in SOD1$^{G93A}$ astrocytes after anti-EMMPRIN antibody treatment. Table referrers to Figure 6.
